# Chromosome segregation is driven by joint microtubule sliding action of kinesins KIF4A and EG5

**DOI:** 10.1101/863381

**Authors:** Kruno Vukušić, Renata Buđa, Ivana Ponjavić, Patrik Risteski, Iva M. Tolić

## Abstract

Successful cell division requires proper chromosome segregation during anaphase. Forces required for chromosome segregation in human cells are linked to sliding of antiparallel microtubules and sliding capacity has been demonstrated *in vitro* for multiple motor proteins, but the molecular mechanism of sliding in the spindle of human cells remains unknown. Using combined depletion and inactivation assays to explore redundancy between multiple targets together with CRISPR technology, we found that PRC1-dependent motor KIF4A/kinesin-4, together with EG5/kinesin-5 motor is essential for spindle elongation in human cells. Photoactivation of tubulin and super-resolution microscopy show that perturbation of both proteins impairs sliding, while decreased midzone microtubule stability cannot explain the observed anaphase arrest. Thus, two independent sliding modules power sliding mechanism that drives spindle elongation in human cells.

---

During anaphase, the conclusive step of mitosis, sister chromatids separate into two daughter cells by kinetochore fiber shortening and spindle elongation (*1–3*). The origin of forces driving spindle elongation in human cells was recently linked to the interpolar region of the spindle (Fig. 1A) where the sliding of antiparallel microtubules (MTs) has been observed (*4*). Although forces required for spindle elongation can be generated at the cell cortex, this is an unlikely scenario in human cells given that depletion of dynein adaptors on the cortex does not impact chromosome segregation during early anaphase (*5*). The sliding mechanism has been proposed years ago (*6*), and *in vitro* experiments demonstrated sliding capacity of multiple motors (*7–10*) but which proteins drive sliding of MTs in mitotic spindle and what is exact molecular mechanism of spindle elongation in human cells remains a longstanding open question.

**Fig. 1.**
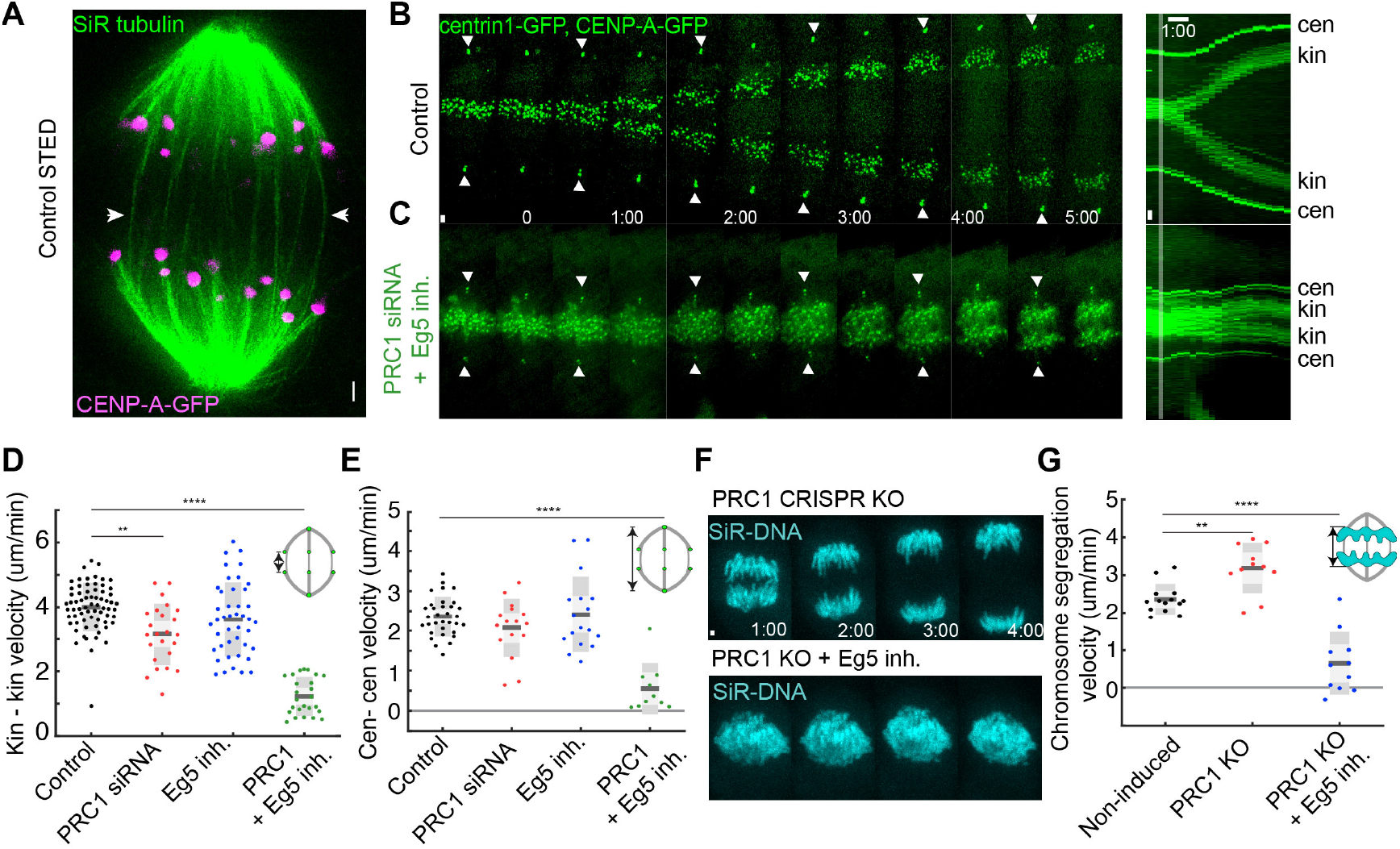
Loss of PRC1 and inactivation of EG5 blocks spindle elongation during anaphase. (**A**) STED image (single z-plane) of anaphase spindle in a live U2OS cell expressing CENP-A-GFP (magenta). Microtubules are labelled with silicon SiR-tubulin (green). Arrowheads point to the spindle midzone where antiparallel microtubules are located. (**B, C**) Live cell images of control and protein regulator of cytokinesis 1 (PRC1) siRNA depleted and S-trityl-L-cysteine (STLC)-treated RPE-1 cell stably expressing CENP-A-GFP and centrin1-GFP. kin-kinetochore and cen-centrosome. White arrowheads indicate centrioles. (**D, E**) Quantification (univariate scatter plot) of velocity of separation of sister kinetochores and spindle elongation velocity (see schemes). Boxes represent standard deviation (dark gray), 95% standard error of the mean (light gray) and mean value (black). Statistics: t test (**P < 0.01, ****P < 0.0001). (**F**) Live cell images of induced RPE-1 PRC1 CRISPR knock-out (KO) and PRC1 KO treated with STLC. Silicon rhodamine (SiR)-DNA was used for chromosome staining. (**G**) Quantification of chromosome segregation velocity (see scheme) in CRISPR experiments. Statistics: t test (**P < 0.01; ****P < 0.0001). Time shown as minutes:seconds. Images are maximum projection of acquired z-stack. Time 0 represents anaphase onset. Vertical scale bars, 1 μm. Horizontal scale bar, 1min.

While spindle elongation can be specifically altered by changing temperature (*11*), global MT dynamics (*12*) or interfering with top signalling effectors (*13*), the aim of this study was to find the force-producing proteins whose depletion stops spindle elongation during anaphase. Although plus-end directed motor kinesin-5 has been implicated as a major candidate for spindle elongation in budding yeast (*14*), its inhibition by a small molecule drug S-trityl-L-cysteine (STLC)(*15*) in RPE-1 human cells, with or without depletion of kinesin-12 (KIF15)(*16*) with a small interfering RNA (siRNA), shortened metaphase spindle, but after anaphase onset did not perturb spindle elongation velocities (fig. S1, A to D and fig. S2, A to I). A significant fraction of cells treated with STLC in metaphase collapsed into monopolar spindles (fig. S1C and fig. S2A), as reported previously (*17*), meaning that EG5-outward force generation is crucial during metaphase, but dispensable during anaphase when inward-forces acting on poles are turned down (*18*). This suggests redundant pathways generating outward forces during anaphase in human cells. Alternatively, force could be generated in the interpolar region by a plus-end directed motor other than EG5. To test this, we depleted the main crosslinker of antiparallel MTs and scaffold for recruitment of multiple motors, protein regulator of cytokinesis 1 (PRC1)(*19*), with a siRNA. PRC1 depletion often led to excessive spindle and cell elongation, as reported previously (*20*), while not affecting rates of spindle elongation in early anaphase (fig. S3, A to E). Accordingly, individual depletions of all PRC1-interacting motor proteins, KIF4A/Kinesin-4 (*21*), KIF23/MKLP-1/Kinesin-6 (fig. S4, A and B), KIF20A/MKLP-2/Kinesin-6 (*22*), centromere-associated protein E (CENP-E)/Kinesin-7 and KIF14/Kinesin-3 (18), did not strongly impact spindle elongation velocities (fig. S5, A to J).

To explore for possible redundancy between EG5 and PRC1-dependent motors we decided to deplete PRC1 along with EG5 inhibition at metaphase-to-anaphase transition, which surprisingly resulted in impaired chromosome segregation by blocking spindle elongation, and cell arrest in early anaphase (Fig. 1, B to E, and fig. S6A), regardless if STLC was added in metaphase or early anaphase (fig. S6B). The poleward motion of chromosomes started but their velocity was slower when compared to controls (fig. S6C). Similar response was observed after inducible knock-out of PRC1 by CRISPR/Cas9 system combined with EG5 inhibition (Fig. 1, F and G, and fig. S7, A to D). Thus, the redundant activity of independent EG5 and PRC1 protein modules (fig. S8, A and B) is crucial for spindle elongation in human cells. Moreover, since PRC1 is a passive MT bundler, we hypothesized that its necessity is due to recruitment of active motor proteins involved in spindle elongation, rather than active sliding.

To test this idea, we depleted PRC1 interacting partners, one by one, in combination with EG5 inhibition. Surprisingly, depletion of KIF4A together with EG5 inhibition completely mirrored the anaphase arrest seen after PRC1 depletion and EG5 inhibition with cells lacking spindle elongation (Fig. 2, A and B, and D and E and fig. S9, A to E), regardless if STLC was added before or after anaphase onset (fig. S9, B to D). Among other candidates, MKLP1, MKLP2, CENP-E and KIF14 were not able to mimic the effect observed after attenuation of EG5 activity and PRC1 depletion (fig. S10, A to J). On the other hand, RNAi mediated co-depletion of MKLP1 and MKLP2 together with EG5 inhibition was reminiscent of EG5 inhibition and KIF4A depletion with cells arrested in early anaphase lacking spindle elongation (Fig. 2, C to E and fig. S11, A to C). This suggests that both human kinesins-6 are required for anaphase spindle elongation, most probably as regulators of KIF4A in the spindle midzone. Therefore, we suggested that MKLP1 and MKLP2 cooperatively regulate KIF4A in the spindle midzone, probably through Aurora B mediated pathway (*23–26*), which then, together with EG5, drive anaphase spindle elongation in human cells. Surprisingly, inhibition of Aurora kinases at anaphase onset also blocked anaphase spindle elongation (fig. S12, A to C), as reported previously (*11*), and co-inhibition of Aurora kinases and EG5 obtained similar results (fig. S12, A to C), suggesting Aurora kinases might separately regulate both sliding modules acting as a major regulators of spindle elongation in human cells.

**Fig. 2.**
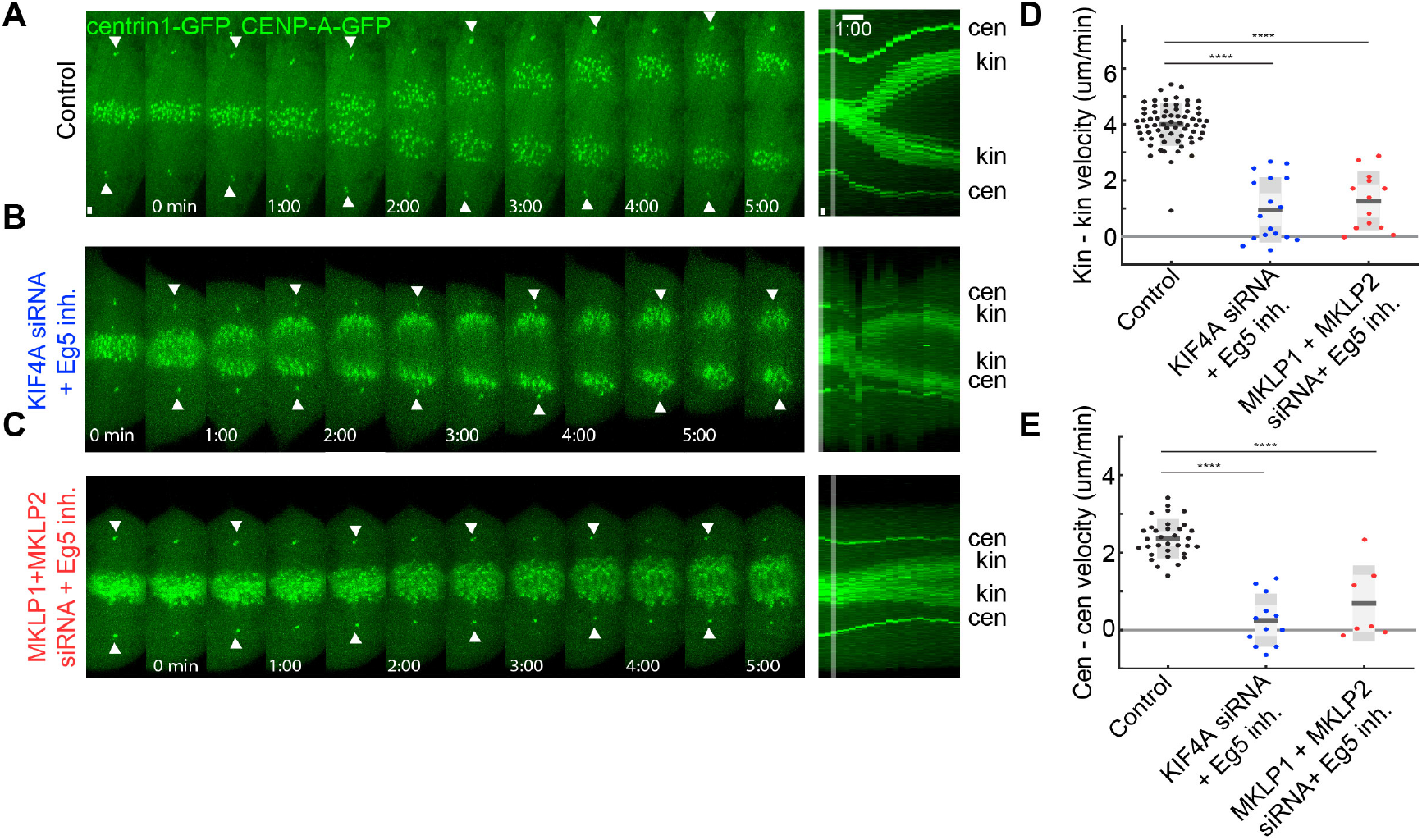
Loss of KIF4A or both kinesins-6 with inactivation of EG5 induces anaphase arrest by blocking spindle elongation. (**A, B, C**) Live cell images of control, KIF4A siRNA depleted and STLC-treated and mitotic kinesin-like protein 1 and 2 (MKLP1 and MKLP2) siRNA depleted and STLC-treated RPE-1 cells stably expressing CENP-A-GFP and centrin1-GFP. kin-kinetochore and cen-centrosome. White arrowheads indicate centrioles. (**D, E**) Quantification of velocity of separation of sister kinetochores and spindle elongation velocity. Statistics: t test (****P < 0.0001). Time shown as minutes:seconds. Images are maximum projection of acquired z-stack. Time 0 represents anaphase onset. Vertical scale bars, 1 μm. Horizontal scale bar, 1min.

To investigate whether lack of midzone MTs (*27*) could explain the observed anaphase arrest we immunolabeled α-tubulin, using a MT-preserving protocol (see Methods), after different treatments and showed that midzone MT organization is similar in control and across the treatments (Fig. 3A, and fig. S13A). Thus, global changes in midzone MT organization cannot explain the anaphase arrest phenotype. Similar results were obtained using super-resolution expansion microscopy (*28*) when comparing EG5-inhibited, KIF4A depleted and EG5-inhibited and control cells (Fig. 3B, and fig. S13B) and stimulated emission depletion (STED) microscopy when comparing induced PRC1 CRISPR/Cas9 knock-out with non-induced control cells (Fig. 3C and fig. S13C). Furthermore, given that depletions of PRC1 or MKLP1 can affect MT stability (*4, 29*) and over-bundling of midzone MTs by means of overexpressing PRC1 can reduce spindle elongation velocity (fig. S14, A to D)(*13, 29*), we decided to investigate whether the arrest phenotype of afore mentioned RNA interference (RNAi) combinations is due to disrupted MT stability. To do so, we used fluorescence dissipation after photoactivation (FDAPA) of photoactivatable (PA)-GFP-α-tubulin to measure MT stability in various conditions (fig. S15). MT stability was reduced significantly only in combinations including depletion of PRC1, as shown previously (*12*), while in the remaining conditions MT stability was not greatly impaired (Fig. 3D and fig. S15). Similar results were obtained using STED microscopy where reduced signal of SiR-tubulin in the midzone was obtained after CRISPR PRC1 knock-out when compared to non-induced controls (fig. S13C). Moreover, we tested whether chromosome segregation and spindle elongation velocities depend on the MT stability across the treatments. Surprisingly, these velocities were not correlated with MT stability (Fig. 3, D and E) demonstrating that the effect of modulating kinesin and PRC1 does not depend on the MT turnover. Spindle elongation during anaphase is thus, contrary to earlier suggestions (*20*), independent of midzone MT stability provided primarily by MT bundling protein PRC1.

**Fig. 3.**
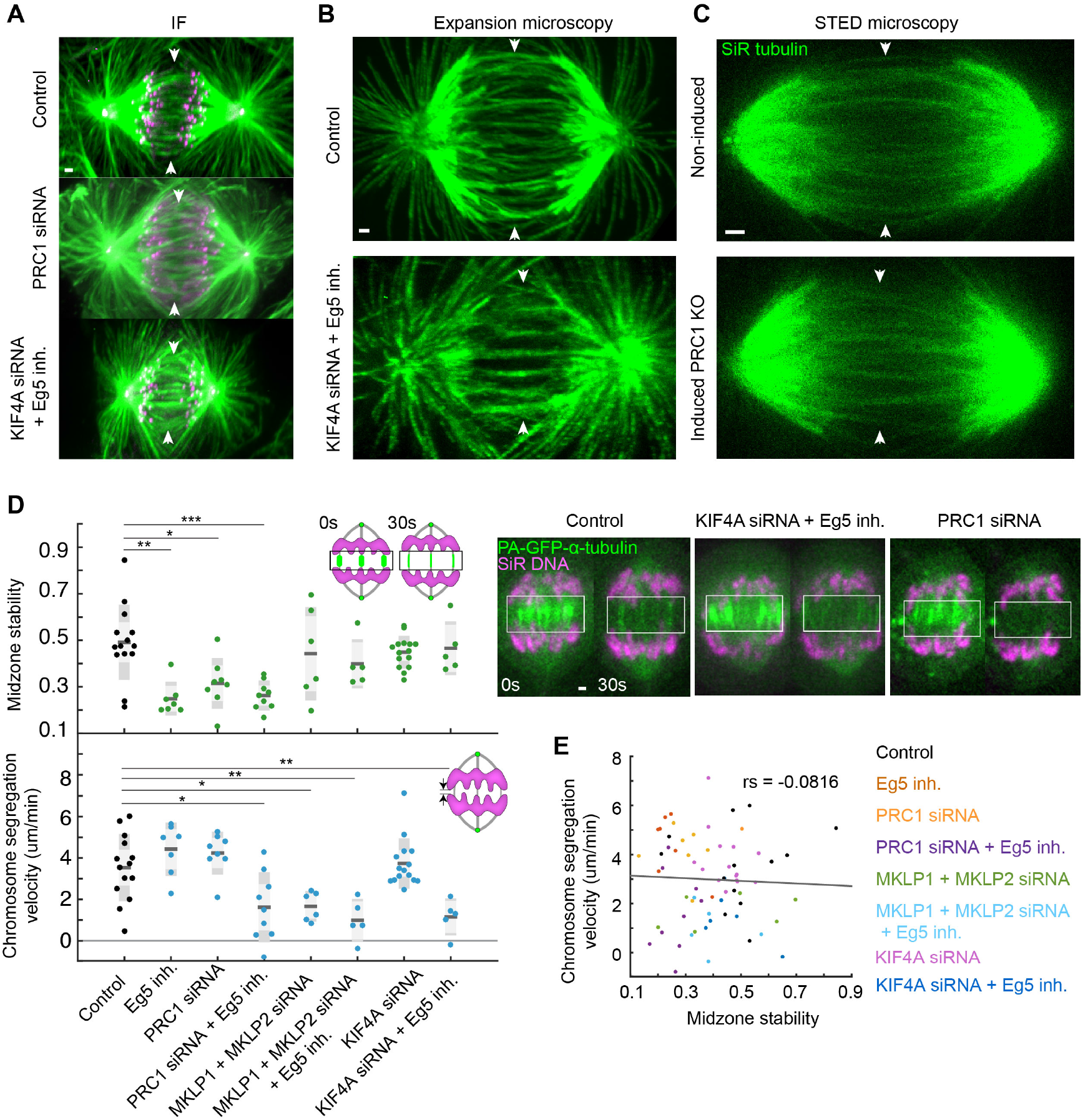
Decreased midzone stability does not affect chromosome segregation velocities during anaphase. (**A**) Immunofluorescent (IF) images of fixed control, PRC1 siRNA-depleted and KIf4A siRNA-depleted and STLC-treated RPE-1 cells stably expressing CENP-A-GFP and centrin1-GFP (magenta) and stained with AlexaFluor594 conjugated with α-tubulin antibody (green). (**B**) Expansion microscopy images of fixed control and KIf4A siRNA-depleted and STLC-treated RPE-1 cells stained with AlexaFluor594 conjugated with α-tubulin antibody. Expansion factor is estimated from spindle length to be 2.3x. (**C**) STED images (single z-plane) of anaphase spindles in a live non-induced control CRISPR PRC1 knock-out (KO) and doxycycline induced PRC1 KO RPE-1 cells. Microtubules are labelled with SiR-tubulin. Arrowheads point to the spindle midzone region. Images are maximum projection of acquired z-stack. **(D)** Smoothed live cell images of RPE-1 cells after photoactivation of photoactivatable (PA)-GFP-α-tubulin in indicated conditions. SiR-DNA (magenta) was used for chromosome staining. Time 0 represents photoactivation onset. Integrated area of intensity was measured using region indicated with a box, along short axis of a box (right). Quantification of ratio of integrated photoactivated signal at time 30s and 0s from onset of photoactivation and chromosome segregation velocity in same time frame in indicated treatments (see schemes). Statistics: t test (*P < 0.05; ** P < 0.01; ***P < 0.001) (left). (**E**) Linear regression and distribution of chromosome segregation velocity versus midzone stability in the different conditions. rs, Spearman correlation coefficient, P < 0.001. Scale bars, 1 μm.

To investigate whether EG5 and KIF4A exert anaphase arrest through antiparallel MT sliding, we measured sliding of midzone MTs after photoactivation of PA-GFP-α-tubulin (fig. S16). The sliding velocity was significantly reduced compared to controls after simultaneous depletion of KIF4A and EG5 inhibition (Fig. 4, A and B, and fig. S16, and fig. S17, A and B). In line with this data, all other conditions impacting KIF4A activity, combined with EG5 inhibition, mimicked this phenotype strongly affecting sliding velocities (Fig. 4B, and fig. S16, and fig. S17, A and B). Moreover, chromosome segregation and spindle elongation velocities strongly correlated with sliding rates across conditions (Fig. 4, B to D, and fig. S17, C and D), which suggests that the origin of anaphase arrest is a result of defective MT sliding. Taken together, KIF4A and EG5 are major plus-end directed sliding motors in anaphase crucial for faithful chromosome segregation in human cells.

**Fig. 4.**
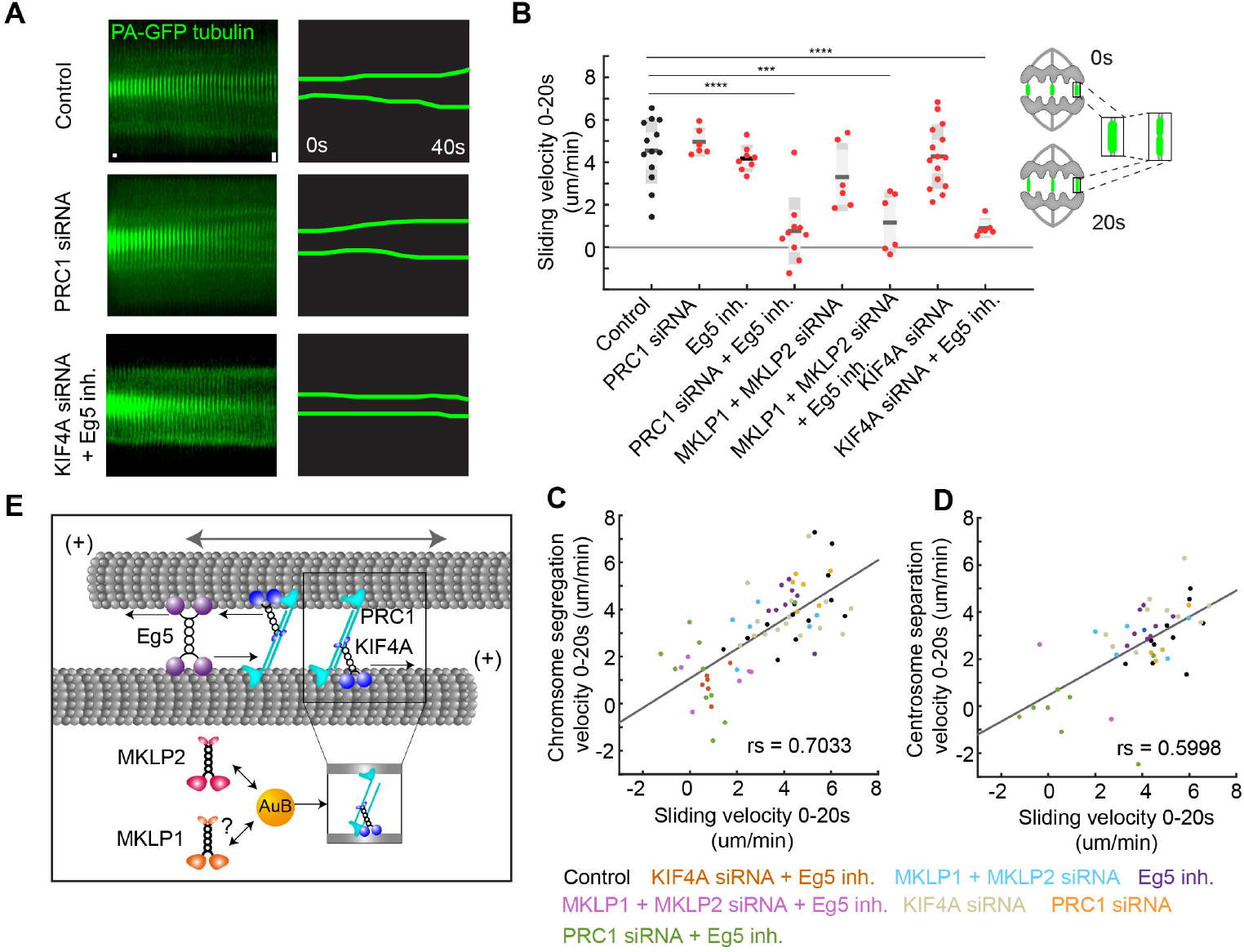
Chromosome segregation and spindle elongation are dependent upon KIF4A and EG5-generated sliding. (**A**) Montage time-lapse live images of mitotic spindle midzone region after photoactivation of photoactivatable (PA)-GFP-α-tubulin in indicated treatments. Note that photoactivated spot does not spread in time after KIf4A siRNA depletion and STLC treatment compared with control and PRC1 siRNA depletion (see scheme generated from kymographs on the left). Horizontal scale bar, 0.8s. Vertical scale bar, 1 μm. (**B**) Quantification of velocity of photoactivated spot spreading (sliding velocity, see scheme) in the different conditions. Statistics: t test (***P < 0.001; ****P < 0.0001). (**C, D**) Linear regression and distribution of chromosome segregation velocity and centrosome separation velocity versus sliding velocity in the different conditions. rs, Spearman correlation coefficient, P < 0.001. (**E**) Proposed model for the sliding in the antiparallel region of anaphase mitotic spindle including two independent sliding modules. One module is composed of plus-end directed motor tetramer EG5 and other is composed of PRC1 dependent plus-end directed motor KIF4A. The latter is probably regulated by Aurora B kinase (AuB) whose midzone localization is dependent on both kinesins-6, MKLP1 and MKLP2. Gray double-headed arrow point to direction of overall microtubule motion sliding over each other as a result of forces produced by microtubule walking on antiparallel microtubules (black arrows).

In summary, our findings answer a long-standing question in cell division. It was unclear what motor proteins drive chromosome segregation and spindle elongation in human cells through sliding mechanism (*3*). Our data reveal that sliding is operated through two independent modules, one composed of EG5 motor protein which generates outward forces throughout cell division (*30*) and the other composed of KIF4A which turns on after anaphase onset (*21*) when KIF4A is brought to midzone MTs by its interaction with PRC1 and is then regulated by a cascade involving Kinesins-6 and Aurora B (Fig. 4E). Sliding forces generated in the midzone antiparallel region by these modules are then transmitted to spindle poles most probably through their lateral connections to k-fibers (*4, 31*). Finally, it would be interesting to study the details of signalling cascade that regulates spindle elongation in human cells where Aurora kinases are promising candidates that could separately regulate both sliding modules (*30, 32, 33*).

## ACKNOWLEDGMENTS

We thank colleagues for providing cell lines and reagents and Andreas Thomae (Core Facility Bioimaging at the Biomedical Center, Ludwig Maximilian University of Munich, Germany) for help with STED microscopy. This work was funded by the European Research Council (ERC Consolidator Grant, GA number 647077), Croatian Science Foundation (HRZZ, project IP-2014-09-4753) and QuantiXLie Centre of Excellence, a project cofinanced by the Croatian Government and European Union through the European Regional Development Fund - the Competitiveness and Cohesion Operational Programme (Grant KK.01.1.1.01.0004). All data is available in the main text or the supplementary materials. The authors declare no conflict of interest.

