## Supplementary Materials for "Chromosome segregation is driven by joint microtubule sliding action of kinesins KIF4A and EG5"

**This PDF file includes:**

Materials and Methods
Supplementary Text
Figs. S1 to S17
Captions for Data S1 to S17
References

### Materials and methods

Cell Lines

The cell lines used are: human U2OS cell lines (human osteosarcoma, female) both wild type and permanently transfected and stabilized using CENP-A-GFP (protein of kinetochore complex), were a gift from Marin Barišić and Helder Maiato (Institute for Molecular Cell Biology, University of Porto, Portugal) (developed in (*1*)), human hTERT-RPE-1 (retinal pigmented epithelium, female) permanently transfected and stabilized using CENP-A-GFP and centrin1-GFP (protein of a centrosome complex), which was a gift from Alexey Khodjakov (Wadsworth Center, New York State Department of Health, Albany, NY) (developed in (*2*)), human hTERT-RPE-1 permanently transfected and stabilized using PA-GFP-α-tubulin which was a gift from Patrick Meraldi (Faculty of Medicine, University of Geneva, Switzerland) (developed in (*3*)), human hTERT-RPE-1 inducible CRISPR/Cas9/PRC1-G2.2 knock-out (KO) which was a gift from Iain Cheeseman (Massachusetts Institute of Technology, Cambridge, MA, USA) (developed and detailed in (*4*)), human HeLa cell lines (human adenocarcinoma, female) separately permanently transfected and stabilized using BAC containing EG5-GFP, MKLP1-GFP and KIF4-GFP which were a gift from Ina Poser and Tony Hyman (Max Planck Institute of Molecular Cell Biology and Genetics, Dresden, Germany) (developed in (*5*)). Cells were grown in flasks in Dulbecco's Modified Eagle's medium (DMEM) with Ultraglutamine (1 g/l D-glucose, pyruvate) (Lonza, Basel, Switzerland) supplemented with 10% of heat-inactivated Fetal Bovine Serum (FBS) (Sigma-Aldrich, St Louis, MO, USA), and penicillin/streptomycin solution (Lonza) to a final concentration of 100 I.U./mL penicillin and 100 µg/mL streptomycin. Media was additionally supplemented for selection of some cell lines as follows: 50 µg/ml geneticin G418 (Life Technologies, Waltham, MA, USA) was added in media for various HeLa BAC cell lines described above and 500 µg/ml G418 was added in media for hTert-RPE-1 PA-GFP-α-tubulin cell line. The induction of RPE-1 PRC1 CRISPR cell line was performed using 1 µg/mL doxycycline hyclate (Sigma) in DMEM media at 24 hr intervals for 4 days (with imaging and analysis on the fifth day), unless otherwise indicated. The cells were kept at 37°C and 5% CO2 in a Galaxy 170s humidified incubator (Eppendorf, Hamburg, Germany). All used cell lines were confirmed to be mycoplasma free by montly checks using MycoAlert Mycoplasma Detection Kit (Lonza) and regular checks while imaging with DNA-stains.

Constructs, transfections and RNAi

U2OS cells were transiently transfected by electroporation using Nucleofector Kit R (Lonza, Basel, Switzerland) with the Nucleofector 2b Device (Lonza, Basel, Switzerland), using X-001 program. Transfection protocol provided by the manufacturer was followed. Cells were transfected with mCherry-PRC1 plasmid provided by Casper C. Hoogenraad (Utrecht University, Utrecht, Netherland). $1 \times 10^6$ cells were electroporated with 1.5 µg of plasmid DNA. Transfection of U2OS cells was performed 25–35 h before imaging. For all siRNA treatments, $2 \times 10^5$ or $3 \times 10^5$ cells were seeded and cultured in 1 ml DMEM medium with same supplements (as above) at 37°C and 5% CO2 on 12-well cell culture plates (Greiner). After one-day growth, at □70% confluency cells were transfected with 200 nM (except KIF15 siRNA: 100 nM and PRC1 siRNA 500 nM) raw targeting or non-targeting siRNA constructs diluted in a Opti-MEM medium (Life Technologies, Waltham, MA, USA). Transfection was performed using Lipofectamine RNAiMAX Reagent (Life Techologies) using protocol provided by the manufacturer. After 5h of treatment the medium was changed to regular

DMEM medium described above. After 24h of treatment cells from one well were equally reseeded into glass bottom microwells with 4 compartments (Grainer), used later for imaging. 3h prior to imaging, the medium was replaced with Leibovit's (L-15) CO2-independent medium (Life Technologies), supplemented as above. The cells were imaged always 48 hours after transfection, unless otherwise indicated. The constructs used were as follows: human MKLP-1 siRNA (sc-35936, Santa Cruz Biotechnology), human KIF14 siRNA (sc-60882, Santa Cruz Biotechnology), human KIF4A siRNA (sc-60888, Santa Cruz Biotechnology), human KIF20A siRNA (sc-91657, Santa Cruz Biotechnology), control siRNA (sc-37007, Santa Cruz Biotechnology), human KIF15 siRNA (L-004960-00-0010, Dharmacon), human PRC1 siRNA (L-019491-00-0020, Dharmacon) and control siRNA (D001810-10-05) from Dharmacon. We observed that inhibition of MKLP1 blocked normal progression of cytokinesis that resulted in formation of binucleated cells. We have observed an increase of 81% in the number of binucleated/multinucleated cells in MKLP1 siRNA-treated samples (calculated from 88 cells) after 48h in comparison with the samples treated with control siRNA (calculated from 92 cells), similar to previous observations (*6*). Similarly, we have observed a significant increase in number of multinucleated cells following 48h treatments with MKLP2 and PRC1 siRNA (*7, 8*). Successful siRNA treatment was checked with immunocytochemistry using appropriate primary antibodies against same targets used in RNAi protocol. Cells were fixed using methanol protocol (described above) 48h after transfection with siRNA and imaged using protocol for imaging of fixed cells (described above).

Drugs

The stock solution of STLC was prepared in dimethyl sulfoxide (DMSO) to a final concentration of 25 mM. Drug was obtained from Sigma-Aldrich. The working solution was prepared in DMEM at 80 μM (the half-maximal inhibitory concentration for STLC in HeLa cells is 700 nM) (*9*). At the time of treatment, the working solution was added to cells at 1:1 volume ratio to obtain a final concentration of 40 μM. To inhibit EG5 (kinesin-5), STLC was added in metaphase or early anaphase. Quick response was observed as most metaphase spindles collapsed into monopolar spindles in RPE-1 cells, minutes after STLC was added (*10*). For immunofluorescence and expansion microscopy of alpha-Tubulin, in treatments with STLC, drug was added to the cell culture media 5 min before fixation. The stock solution of GSK-923295 was prepared in dimethyl sulfoxide (DMSO) to a final concentration of 8 mM. GSK-923295 was obtained from MedChemExpress (MCE, NJ, USA). The working solution was prepared in DMEM at 200 nM. At the time of treatment, the working solution was added to cells at 1:1 volume ratio to obtain a final concentration of 100 nM (IC50 value of compound is 3.2 nM) (*11*). To inhibit CENP-E, GSK-923295 was added in late metaphase. Appearance of spindles blocked in prometaphase with fraction of kinetochores trapped around polar region of the spindle (*12*) confirmed the effect of GSK-923295, imaged 30 min post-treatment with drug. The stock solution of ZM 447439 was prepared in dimethyl sulfoxide (DMSO) to a final concentration of 2 mM. ZM 447439 was obtained from Selleckchem (Munich, Germany). The working solution was prepared in DMEM at 8 μM. At the time of treatment, the working solution was added to cells at 1:1 volume ratio to obtain a final concentration of 4 μM (IC50 value of compound is 110 nM for Aurora A and 130 nM for Aurora B)(*13*). To inhibit Aurora kinases, ZM 447439 was added at metaphase-to-anaphase transition. In experiment where KIF15 was inhibited by siRNA treatment in U2OS cells, we observed rapid collapse of metaphase spindle upon 40 μM STLC treatment, as reported previously (*14*).

Sample preparation

When cells reached 80% confluence, DMEM medium was removed from the flask and the cells were washed with 5 mL of 1% PBS. Afterward, 1 mL of 1% Trypsin/EDTA (Biochrom AG, Berlin, Germany) was added and the cells were incubated at 37 °C and 5% CO2 in a humidified incubator (Eppendorf). After 5 min incubation, Trypsin was blocked by adding 2-5 mL of DMEM medium. Cells were counted using the Improved Neubauer chamber (BRAND GMBH + CO KG, Wertheim, Germany) and $4.5x10^5$ cells were seeded and cultured in 2ml DMEM medium with same supplements (as above) at 37°C and 5% CO2 on 14 or 20 mm glass microwell uncoated 35mm dishes with 0.16-0.19mm (#1.5 coverglass) glass thickness (MatTek Corporation, Ashland, MA, USA). For siRNA experiments, $1x10^5$ cells were seeded in cell culture 35/10 mm glass bottom microwells with 4 compartments (Greiner, Frickenhausen, Germany) or $2x10^5$ in 12-well cell culture plates (Greiner) that were reseeded the day after into glass bottom microwells with 4 compartments. After one-day growth, 3h prior to imaging, the medium was replaced with Leibovit's (L-15) CO2-independent medium (Life Technologies), supplemented with 10% FBS (Life Technologies), 100 I.U./mL penicillin and 100 µg/mL streptomycin. For live-cell staining of chromosomes 1 hour before imaging, silicon rhodamine (SiR)-DNA, also called SiR-Hoechst (*15*) (Spirochrome AG, Stein am Rhein, Switzerland) was added to 1 mL of cells in a DMEM medium to a final concentration of 100 nM together with efflux pump inhibitor verapamil (Spirochrome AG), only in RPE-1 and U2OS cell lines, to a final concentration of 10 µM.

Live cell imaging

HeLa cells expressing EG5-GFP were imaged using a Leica TCS SP8 X laser scanning confocal microscope with a HC PL APO ×63/1.4 oil immersion objective (Leica, Wetzlar, Germany) heated with an objective integrated heater system system (Okolab, Pozzuoli, NA, Italy). For excitation, a 488-nm line of a visible gas Argon laser and a visible white light laser at 575 nm were used for GFP and mRFP, respectively. GFP and mRFP emissions were detected with HyD (hybrid) detectors in ranges of 498–558 and 585–665 nm, respectively. Pinhole diameter was set to 0.8 µm. Images were acquired at 30–60 focal planes with 0.5 µm z-spacing, 30 nm xy-pixel size, and 400 Hz unidirectional xyz scan mode. The system was controlled with the Leica Application Suite X Software (1.8.1.13759, Leica, Wetzlar, Germany) and cells were maintained at 37 °C and in 5% CO2 using Okolab stage top heating chamber (Okolab, Pozzuoli, NA, Italy). All RPE-1 and U2OS cells were imaged using Bruker Opterra Multipoint Scanning Confocal Microscope (detailed in (*16*)) (Bruker Nano Surfaces, Middleton, WI, USA). The system was mounted on a Nikon Ti-E inverted microscope equipped with a Nikon CFI Plan Apo VC ×100/1.4 numerical aperture oil objective (Nikon, Tokyo, Japan). During imaging, cells were maintained at 37 °C and 5% CO2 in Okolab Cage Incubator (Okolab, Pozzuoli, NA, Italy). A 60 µm pinhole aperture was used and the *xy*-pixel size was 83 nm. For excitation of DAPI, GFP, mCherry or RFPa and SiR fluorescence, a 405, 488, 561 and 647 nm diode laser line was used, respectively. The excitation light was separated from the emitted fluorescence by using Opterra Dichroic and Barrier Filter Set 405/488/561/640 nm (DAPI/eGFP/TRITC/Cy5) (Chroma, USA). The brightfield imaging of RPE-1 PRC1 KO cells in different conditions was performed by illuminating sample with brightfield lamp (Nikon). Images were captured with an Evolve 512 Delta EMCCD Camera using 150 ms exposure time (Photometrics, Tucson, AZ, USA) with no binning performed. In experiments where whole spindle stack was imaged, z-stacks were acquired at 30-60 focal planes separated by 0.5 µm with unidirectional xyz scan mode and with "Fast Acquisition" option in software enabled. Otherwise, one or few z-stacks were imaged using 0.5 µm spacing with unidirectional xyz scan mode. The system was controlled with the Prairie View Imaging Software (Bruker Nano Surfaces, Middleton, WI, USA).

Photoactivation (PA) stability and sliding assays

For photoactivation of fluorescence of PA-GFP, a 405-nm laser diode (Coherent, Santa Clara, CA, USA) was used. Photoactivation was performed using live photoactivation option in PraireView software, with duration of pulse set to 400 ms for each point and laser power set to 40% for all experiments performed. Photoactivation was performed in a line pattern on an equally distributed 10 points, where each point represents one laser hit. The interval between the points was minimal, 0.05 ms, and photoactivation area was set to 0.5 μm for each point. Photoactivation was performed after anaphase onset, perpendicular to spindle long exis, in between separating chromatids, visualized with 100 nM SiR-DNA stain and detected using maiximum power of 0.5% of 647 diode laser. SiR-DNA was used to discriminate anaphase onset by perception of peripheral sister chromatids moving into characteristic anaphase "v-shape", which was then used as a mark for start of the photoactivation assay. PA-GFP fluorescence was detected using 488 diode laser (on 30% of a maximum laser power), turned on just after onset of anaphase to minimalize unwanted photoactivation. The excitation light was separated from the emitted fluorescence in both channels by using Opterra Dichroic and Barrier Filter Set 488/640 nm (eGFP/Cy5) (Chroma, USA). The interval between consecutive frames was set to 0.8 s, imaging one central z-plane and program was set to record 200 consecutive frames. Further details of this microscopy system and similar laser ablation and photoactivation assays were discussed elsewhere (*16*).

Immunofluorescence

Human RPE-1 cell line stably expressing centrin1-GFP and CENP-A-GFP were grown on glass-bottomed dishes (14 mm, No. 1.5, MatTek Corporation) and fixed by 1 mL of ice-cold methanol for 3 min at -20°C for visualization of PRC1, EG5, KIF15, KIF4A, MKLP1, MKLP2 and KIF14. To visualize alpha-Tubulin in control cells and in the treatments, ice-cold methanol protocol was avoided because it destroys unstable microtubules and cells were instead fixed by a mixture of 3.2% PFA (paraformaldehyde) and 0.25% GA (glutaraldehyde) in microtubule-stabilizing PEM buffer (0.1 M PIPES, 0.001 M $MgCl_2$ x 6 $H_2O$, 0.001 M EDTA, 0.5 % Triton-X-100) for 10 min at room temperature (*17*). After fixation with PFA and GA, for quenching, cells were incubated in 1mL of freshly prepared 0.1% borohydride in PBS (phosphate-buffered saline) for 7 min and after that in 1 mL of 100 mM $NH_4Cl$ and 100 mM glycine in PBS for 10 min at room temperature. Both methanol fixed cells and PFA and GA fixed cells were then washed with 1 mL of PBS, 3 times for 5 min. To block unspecific binding of antibodies, cells were incubated in 500 μL blocking/permeabilization buffer (2% normal goat serum (NGS) and 0.5% Triton-X-100 in water) for 45 min at room temperature. Cells were then incubated in 500 μl of primary antibody solution for 24h at 4 °C. The following primary antibodies were used: mouse monoclonal PRC1 (sc-376983, Santa Cruz Biotechnology), diluted 1:50; mouse monoclonal EG5 (sc-365681, Santa Cruz Biotechnology), diluted 1:50; mouse monoclonal KIF15 (sc-100948, Santa Cruz Biotechnology), diluted 1:50; mouse monoclonal KIF4A (sc-365144, Santa Cruz Biotechnology), diluted 1:50; rabbit polyclonal MKLP-1 (sc-867, Santa Cruz Biotechnology), diluted 1:50; mouse monoclonal KIF20A (sc-374508, Santa Cruz Biotechnology), diluted 1:50;, rat anti-alpha Tubulin (Invitrogen), diluted 1:500. After primary antibody, cells were washed in PBS and then incubated in 500 μL of secondary antibody solution for 45 min at room temperature. Alexa Fluor 488, 594, and 647 (Invitrogen) were used as secondary antibodies (diluted 1:1000).

Expansion microscopy

Expansion microscopy protocol was a custom made as a combination of steps from of various previously developed protocols for expansion microscopy of human cells in culture (*18-20*). Human RPE-1 cell line stably expressing centrin1-GFP and CENP-A-GFP were grown on glass-bottomed dishes (14 mm, No. 1.5, MatTek Corporation) and fixed by a mixture of 3.2 % PFA (paraformaldehyde) and 0.25 % GA (glutaraldehyde) in PEM buffer (0.1 M PIPES, 0.001 M $MgCl_2$ x 6 $H_2O$, 0.001 M EDTA, 0.5 % Triton-X-100) for 10 min at room temperature. After fixation with PFA and GA, for quenching, cells were incubated in 1mL of freshly prepared 0.1% borohydride in PBS (phosphate-buffered saline) for 7 min and after that in 1 mL of 100 mM $NH_4Cl$ and 100 mM glycine in PBS for 10 min at room temperature. Cells were then washed with 1 mL of PBS, 3 times for 5 min. To block unspecific binding of antibodies, cells were incubated in 500 µl blocking/permeabilization buffer (2% normal goat serum (NGS) and 0.5% Triton-X-100 in water) for 45 min at room temperature. Cells were then incubated in 500 µl of primary antibody solution for 24h at 4 °C. Primary antibody that was used is rat anti-alpha Tubulin (Invitrogen). After primary antibody, cells were washed in PBS and then incubated in 500 µl of secondary antibody solution for 45 min at room temperature. Secondary antibody is Alexa Fluor 594 (Invitrogen). To remove unbound secondary antibody from the sample, cells were washed three times with 1 ml of PBS for 5 min each, at room temperature. Acryloyl-X 1:100 (vol/vol) (Thermo Fisher Scientific) was diluted to 0.1 mg/ml in anchoring buffer. Sample was incubated with the anchoring solution for at least 6 h or overnight. Wash the sample two times in 1 ml of PBS, 5 min each, at room temperature, right before application of the gelation solution. Mix the gel monomer components (8.6 wt% Sodium acrylate, 2.5 wt% Acrylamide, 0.15 wt% N,N'-Methylenebisacrylamide, 11.7 wt% Sodium chloride, 1x PBS and water). Apply the polymerization solution to the sample and let the gel polymerize by incubating for 1 h in a humidified chamber, at 37 °C. After that, incubate sample in digestion buffer (50 mM Tris pH 8.0, 1 mM EDTA, 0.5% Triton-X-100, 0.8 M guanidine HCl) with proteinase K (1:100, final concentration 8 units/mL) for >8 h, or overnight in a humidified digestion chamber, at room temperature. After digestion, remove the remaining digestion buffer from the dish and add at least 2 mL $ddH_2O$. Remove the $ddH_2O$ after 10–20 min of incubation, using a Pipetboy or a vacuum pump. Add fresh $ddH_2O$. Repeat this step until no further expansion of the gel can be observed (usually after four to five water exchanges). Before the imaging, excess water was removed by pipette and after that by placing filter paper in the corners of the dish to minimalize movement of the gel during imaging. Expansion factor is estimated from measured spindle length of the expanded sample after dividing it with the spindle length of the non-expanded spindle in the same phase of the anaphase.

Imaging of fixed cells

All RPE-1 and U2OS cells fixed cells were imaged using Bruker Opterra Multipoint Scanning Confocal Microscope (Bruker) described above. In experiments where whole spindle stack was imaged, z-stacks were acquired at 30-60 focal planes for immunofluorescence images, and 60-120 focal planes for expanded samples, separated by 0.5 µm with unidirectional xyz scan mode. A 60 µm pinhole aperture was used and the *xy*-pixel size was 83 nm. For excitation of DAPI, GFP, mCherry or RFPa and SiR fluorescence, a 405, 488, 561 and 647 nm diode laser line was used, respectively. The excitation light was separated from the emitted fluorescence by using Opterra Dichroic and Barrier Filter Set 405/488/561/640 nm (DAPI/eGFP/TRITC/Cy5) (Chroma). Images were captured with an Evolve 512 Delta EMCCD Camera using 300 ms exposure time (Photometrics, Tucson, AZ, USA) with no binning performed. The frame average was performed 8 times for imunoflurescence images and 16 times for expansion microscopy images. All experiments were carried out using Nikon CFI Plan Apo VC ×100/1.4 numerical aperture oil objective (Nikon).

STED microscopy

STED images of U2OS and RPE-1 cells were recorded at the Core Facility Bioimaging at the Biomedical Center, LMU Munich. STED resolution images were taken of SiR-tubulin signal, whereas GFP signal of kinetochores was taken at confocal resolution. Gated STED images were acquired with a Leica TCS SP8 STED 3X microscope with pulsed white light laser excitation at 652 nm and pulsed depletion with a 775 nm laser (Leica, Wetzlar, Germany). The objective used was HC PL APO CS2 ×93/1.30 GLYC with a motorized correction collar set to 63%. Scanning was done at 30 Hz, a pinhole setting of 0.93 AU (at 580 nm), and the pixel size was set to 33.29 × 33.29 nm. The signals were detected with Hybrid detectors with the following spectral settings: SiR-tubulin (excitation 652; emission: 662–692 nm; counting mode, gating: 0.35–6 ns) and GFP (excitation 488; emission 498–550; counting mode, gating: 0.50-6 ns). STED 775 nm laser was delayed by -150 ps. Cells were stained with SiR-tubulin dye at 100 nM concentration, 1 h before imaging.

Parameters used to define metaphase-to-anaphase transition

The time of anaphase A onset for each individual cell was defined as the time point immediately prior to the separation of sister chromatid populations as annotated manually using visual inspection based on the increased distance between the sister kinetochore groups as cells transitions from metaphase to anaphase. The onset of spindle elongation was defined as the time point immediately prior to the separation of two centrosomes as annotated by checking for continuous increase in the spindle length for two consecutive frames using custom made Matlab script (The MathWorks Inc., USA, R2018a) and double checked by visual inspection of original movies for possible errors.

Tracking of kinetochore, centrosome and chromosome motions

Kinetochores and centrosomes were tracked in time using Low Light Tracking Tool (LLTT), an ImageJ plugin (*21*). Tracking of kinetochores in the x, y plane was performed on individual imaging planes or on maximum-intensity projections of up to three planes. The position in z direction (3D) was ignored because it had a small contribution to the kinetochore movement. In order to obtain optimal tracking results, it was necessary to define good intensity offset in the channel with fluorescently labeled kinetochores. The intensity offset was defined by measuring the mean intensity around kinetochores in the first frame before start of tracking using 'freehand selection' tool in Fiji. Sometimes, when photobleaching was prominent, bleach correction using Histogram Matching Method in Fiji was done to compensate for a decrease in background intensity in time. Also, it was necessary to define EMCCD gain and Electrons per A/D count of the used EMCCD camera to correct the measured flux of the object and background noise. The EMCCD-GaussianML tracking algorithm method was used (*21*) because it yielded more precise results compared with Gaussian-ML method, especially in situations when fast movement of the tracked object occurred (on a scale of micron per frame or more). All tracked objects were double checked by eye to ensure that tracking was accurate, because it was inaccurate in situations of an uneven intensity of tracked objects and in situations when multiple similar objects appeared in close proximity. If those cases were predominant, tracking was performed manually extracting xyz-coordinates of each kinetochore. σ value (standard deviation of the Gaussian used to approximate the Point Spread Function (PSF) of the tracked objects) was set to 1 to encompass just the tracked kinetochore. Detailed quantitative analysis of centrosome and kinetochore location was performed using custom made MATLAB scripts which determines the distance between the centroid of the sister kinetochores. Chromosomes where tracked using ImageJ line tool from a centromere region of a one sister chromosome to the same region of another sister. The velocities of kinetochore and centrosome separation were measured between 1 and 3 minutes after the onset of anaphase, $(d_3-d_1)/(2t)$ where d is the distance between sister kinetochore centroids subtracted to the same distance in the last frame before anaphase or pole-to pole distance subtracted to the same distance in the last frame before anaphase (termed delta for all parameters defined this way) and t is the time period. This interval is chosen because these parameters are linearly increasing during this time period in untreated human cells (*22, 23*).

Quantification of microtubule stability

Microtubule stability was measured on single z-plains acquired in photoactivation assay. The onset of photoactivation was set as a first point of measurement, after all dots finished on predetermined photoactivation line, and second point of measurement was placed in a frame 30s after onset in all conditions imaged. Region between separated chromosomes in the last frame was used as a borderline for drawing a 100px thick line in ImageJ, horizontal to the spindle long axis, from which the fluorescence profile was extracted. The mean background fluorescent intensity measured in the same frame in cytoplasm (measured in ImageJ by drawing a 100px thick line), was subtracted from obtained intensities. The obtained values were plotted in SciDavis program (Free Software Foundation, Inc., Boston, MA, USA) and area under the peak was linearly integrated to obtain area of intensity for that frame. The same was performed for the frame defined as the onset of photoactivation using region of the same dimensions. The two values were divided to give estimation on how much of photoactivated-a-tubulin fluorescence was lost following 30 s period in all conditions tested giving estimation of relative microtubule turn-over and stability.

Quantification of sliding velocity

The hTERT-RPE-1 permanently transfected and stabilized using PA-GFP-α-tubulin where imaged in a one middle z-plane using unidirectional xyz scan mode and with "Fast Acquisition" option in software enabled with interval of 0.8s between two frames. The width of photoactivated spot in the spindle midzone was measured using ImageJ by drawing a line segment along the photoactivated region, from the moment when photoactivation of tubulin finished for all dots on line segment used for photoactivation, across time frames separated by 0.8s, to the last frame frame of measurable photoactivation signal in the spindle midzone. The width was plotted as a function of time in Matlab home written script and on each curve linear regression was performed for points starting from 0, representing start of a measurement, to the 30s after. The velocities where calculated from a slope of a regression line. The position of a centrosome was determined only in those cells in which signal of photoactivated a-tubulin could be discerned on both poles. Multiple bundles were tracked in single cell were appropriate but their velocities values were averaged for correlation graphs and bar plots.

Quantification of PRC1 overexpression

The fluorescence intensity signal of PRC1-mCherry was measured on the whole spindle region using ImageJ on sum-intensity projection of all z-stacks acquired. The background fluorescence intensity measured in the cytoplasm was subtracted from the mean value obtained, and this value was divided with number of z-stacks used in sum projection.

CRISPR KO cell scoring

CRISPR PRC1 KO cell were scored for successful knockout of PRC1 5dy post-induction with doxycycline by fixing cells with cold-methanol protocol (described above) and staining with mouse monoclonal PRC1 antibody (sc-376983, Santa Cruz Biotechnology), diluted 1:50, and Alexa Fluor 488 (Invitrogen) secondary antibody (diluted 1:1000). SiR-DNA, 100nM final concentration, was added post-fixation, 30 min before imaging, to stain DNA in order to identify anaphase cells. Successful KO without noticeable PRC1 signal in the spindle midzone was observed in 90% of analysed cells (60 out of 68 cells) and 10% of the cells imaged had a normal PRC1 appearance, localising to the spindle midzone, similarly to all non-induced controls imaged.

Image processing and statistical analysis

Image processing was performed in ImageJ (National Institutes of Health, Bethesda, MD, USA). Quantification and statistical analysis were performed in MatLab. Figures were assembled in Adobe Illustrator CS5 and Adobe Photoshop CS5 (Adobe Systems, Mountain View, CA, USA). All kymographs were generated in ImageJ on maximum intensity projections of vertically rotated images by using a "Reslice" command and them performing maximum intensity projection and montage images were generated in ImageJ in single focal plane vertically oriented images using "Make Montake" command. The kymograph and montage images were rotated in every frame to fit the long axis of the spindle to be parallel with the central long axis of the box in ImageJ and spindle short axis to be parallel with the central short axis of the designated box in ImageJ. The designated box sizes were cut in the same dimensions for all panels in Figures where the same experimental setups were used across the treatments. When comparing different treatments in channels with same proteins labelled, minimum and maximum of the intensity in that channel was set to the values in the control treatment. When indicated, smoothing of images was done using Gaussian blur function in ImageJ (s=1.0-1.5). Data are given as mean ± s.t.d., unless otherwise stated. p values were obtained using unpaired two-sample Student's t-test (significance level was 5%). When comparing the same parameters cell by cell, we used paired two-sample Student's t-test (significance level was 5%). $p < 0.05$ was considered statistically significant, very significant if $0.001 < p < 0.01$ and extremely significant if $p < 0.001$. Values of all significant differences are given with degree of significance indicated (*$0.01 < p < 0.05$, **$0.001 < p < 0.01$, ***$p < 0.001$, **** $< 0.0001$). For linear regression correlation measure between two parameters, nonprametric Spearman correlation coefficient, termed rs, was used where $p<0.001$. The number of analysed cells and specific parameters is given in the respective figure panel.

### Supplementary text: Extended acknowledges and author contributions

We thank M. Barišić, H. Maiato, A. Khodjakov, P. Meraldi, I. Cheeseman, C.C. Hoogenraad, I. Poser and T. Hyman for sharing their regents and cell lines; Dora Zvjerković for help in data analysis; Sonja Lesjak for technical help in cell culture maintenance; all members of I.M.T and N. Pavin groups for helpful discussions and/or critical reading of this manuscript; and Ivana Šarić for the drawings. All the experiments were performed by K.V., R.B. and I.P. in the laboratory of I.M.T except STED microscopy and experiments on HeLa cell lines which were performed by P.R.; K.V., R.B. and I.P. analysed the data; K.V. quantified the data; K.V., R.B., I.P. and I.M.T. wrote the manuscript; I.M.T. coordinated the project.

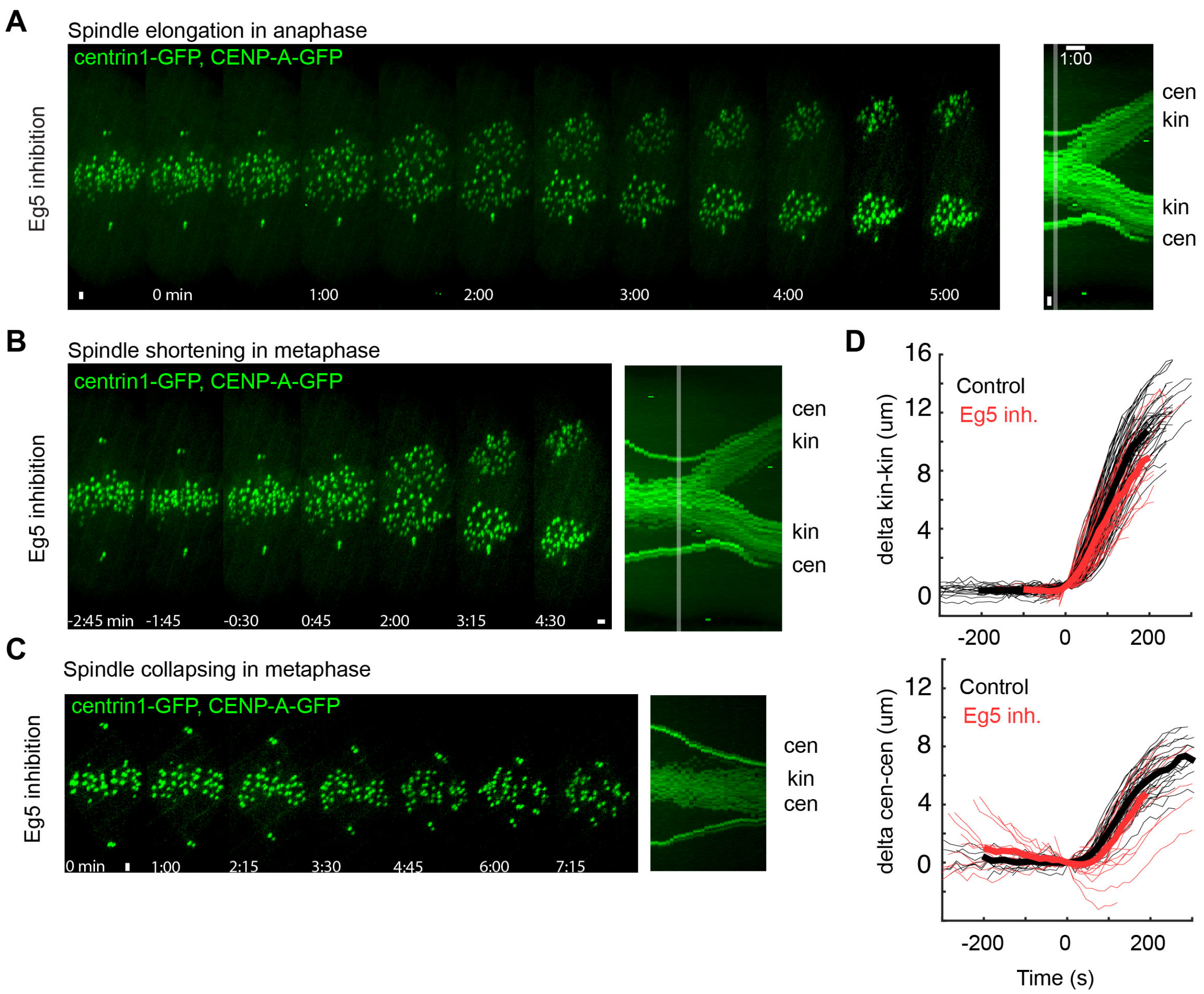


**Fig. S1.** Inhibition of EG5 is crucial for maintenance of metaphase spindle length but dispensable for anaphase spindle elongation. (A) Live cell images of S-trityl-L-cysteine (STLC)-treated RPE-1 cell stably expressing CENP-A-GFP and centrin1-GFP. kin-kinetochore and cen-centrosome. (B) Live cell images of STLC-treated RPE-1 cell show shortening of spindle in metaphase and normal spindle elongation in anaphase. Time 0 represents anaphase onset. (C) Live cell images of RPE-1 cell show spindle collapsing during metaphase after STLC treatment. (D) Plots of relative kinetochore and centrosome separation distance (delta) defined as the kinetochore-to-kinetochore (kin-kin) and centrosome-to-centrosome (cen-cen) distance at time t minus the kin-kin and cen-cen distance at t = 0, over time. Individual kinetochore and centrosome pairs (thin lines), mean (thick lines). Time zero represents start of STLC treatment. Images are maximum projection of acquired z-stack. Time shown as minutes:seconds. Vertical scale bars, 1 μm. Horizontal scale bar, 1min.

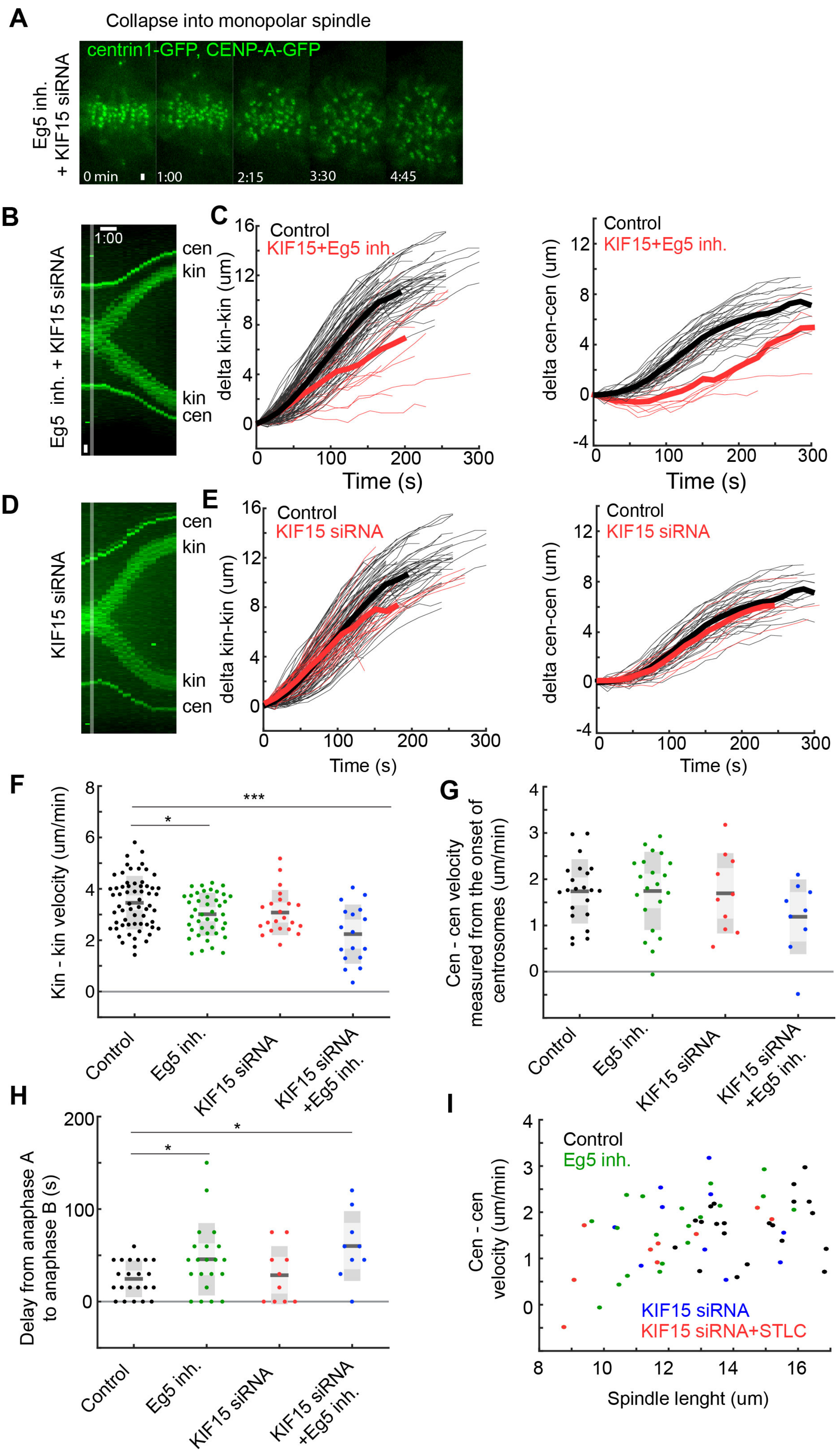
A
Collapse into monopolar spindle
centrin1-GFP, CENP-A-GFP
Eg5 inh. + KIF15 siRNA
0 min
1:00
2:15
3:30
4:45
B
Eg5 inh. + KIF15 siRNA
1:00
cen
kin
kin
cen
C
Control
KIF15+Eg5 inh.
delta kin-kin (um)
Time (s)
delta cen-cen (um)
D
KIF15 siRNA
E
Control
KIF15 siRNA
F
Kin - kin velocity (um/min)
G
Cen - cen velocity measured from the onset of centrosomes (um/min)
Control
Eg5 inh.
KIF15 siRNA
KIF15 siRNA +Eg5 inh.
H
Delay from anaphase A to anaphase B (s)
I
Cen - cen velocity (um/min)
Spindle lenght (um)
KIF15 siRNA
KIF15 siRNA+STLC

**Fig. S2.** Depletion of KIF15 and inhibition of EG5 combined with KIF15 depletion does not affect anaphase spindle elongation. (A) Live cell images of KIF15 siRNA depleted and S-trityl-L-cysteine (STLC)-treated RPE-1 cell stably expressing CENP-A-GFP and centrin1-GFP showing metaphase spindle collapse and monopolar spindle formation. kin-kinetochore and cen-centrosome. Time zero represents start of STLC treatment. (B) Kymograph (consecutive maximal intensity projections onto the x-axis) of the live cell images of KIF15 siRNA depleted and S-trityl-L-cysteine (STLC)-treated RPE-1 cell stably expressing CENP-A-GFP and centrin1-GFP. (C) Plots of relative kinetochore and centrosome separation distance (delta) defined as the kinetochore-to-kinetochore (kin-kin) and centrosome-to-centrosome (cen-cen) distance at time t minus the kin-kin and cen-cen distance at $t = 0$, over time. Individual kinetochore and centrosome pairs (thin lines), mean (thick lines). (D) Kymograph of the live cell images of KIF15 siRNA depleted RPE-1 cell. (E) Plots of relative kinetochore and centrosome separation distance over time. (F) Quantification (univariate scatter plot) of velocity of separation of sister kinetochores measured from the onset of anaphase. Boxes represent standard deviation (dark grey), 95% standard error of the mean (light grey) and mean value (black). Statistics: t test (***$P < 0.001$; ****$P < 0.0001$). (G) Quantification of velocity of separation of spindle elongation measured from the onset of centrosome separation. (H) Quantification of spindle elongation delay (anaphase B) calculated as the difference between spindle elongation onset and kinetochore separation onset (anaphase A). Statistics: t test (*$P < 0.05$). Note: EG5 inhibition by STLC treatment, with or without depletion of KIF15, induces a small delay in spindle elongation start compared to controls. (I) Linear regression and distribution of spindle elongation velocity (cen-cen) versus inital spindle length, measured at the beginning of anaphase, in indicated conditions. Note: initial spindle length and spindle elongation velocities are not correlated across conditions. Images are maximum projection of acquired z-stack. Time shown as minutes:seconds. Time 0 represents anaphase onset. Vertical scale bars, 1 μm. Horizontal scale bar, 1min.

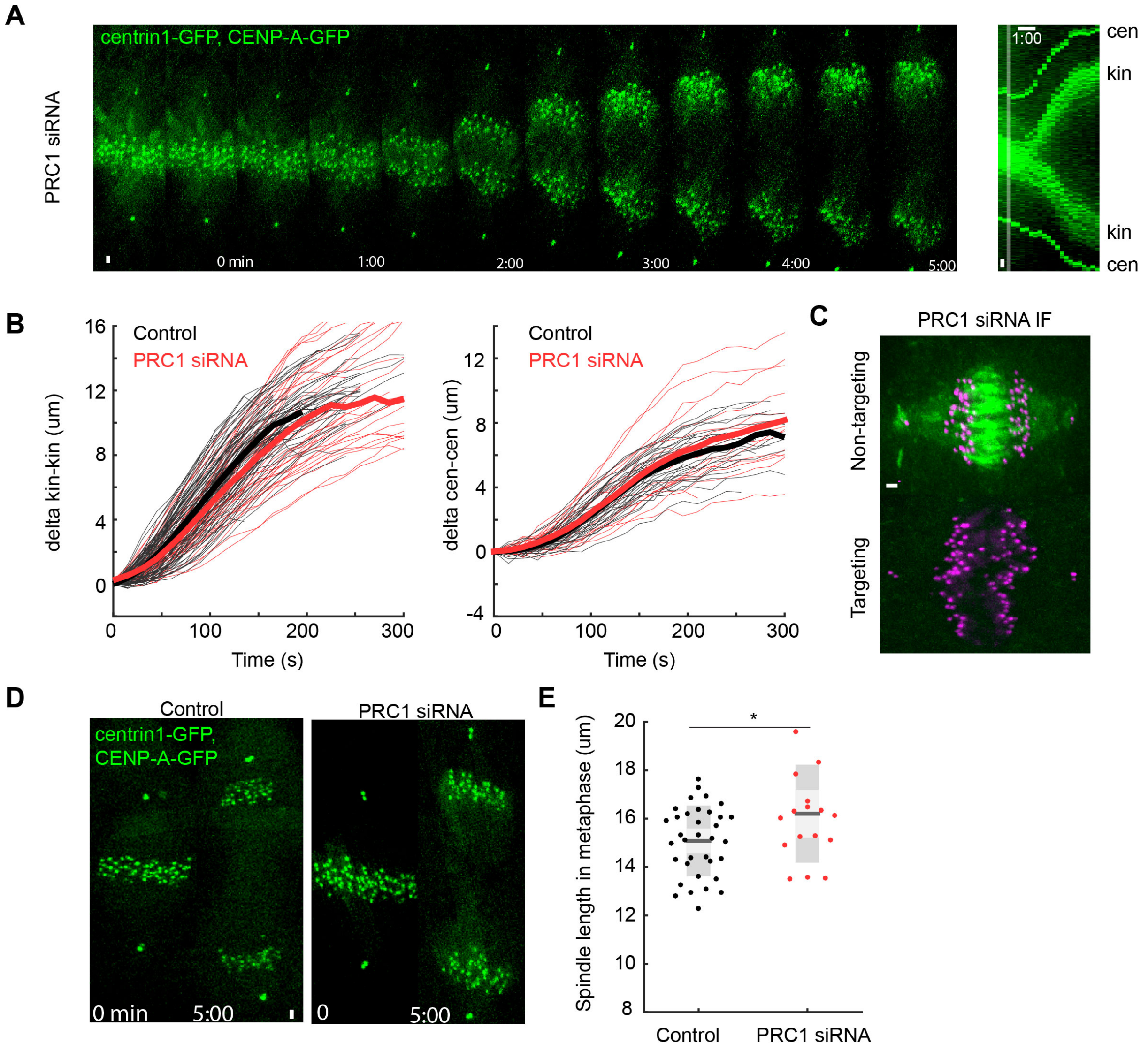


**Fig. S3.** Depletion of PRC1 does not affect anaphase spindle elongation. (A) Live cell images of PRC1 siRNA depleted RPE-1 cell stably expressing CENP-A-GFP and centrin1-GFP show unperturbed spindle elongation. kin-kinetochore and cen-centrosome. (B) Plots of relative kinetochore and centrosome separation distance (delta) defined as the kinetochore-to-kinetochore (kin-kin) and centrosome-to-centrosome (cen-cen) distance at time t minus the kin-kin and cen-cen distance at t = 0, over time. Individual kinetochore and centrosome pairs (thin lines), mean (thick lines). (C) Immunofluorescence (IF) images of fixed non-targeting treated and PRC1 siRNA-depleted RPE-1 cells stably expressing CENP-A-GFP and centrin1-GFP (magenta) stained with AlexaFluor594 conjugated with PRC1 antibody (green). (D) Live cell images of control and PRC1 siRNA depleted RPE-1 cell show longer metaphase and anaphase spindles after PRC1 depletion. (E) Quantification (univariate scatter plot) of spindle length in metaphase in treatments as in (D). Boxes represent standard deviation (dark grey), 95% standard error of the mean (light grey) and mean value (black). Statistics: t test (*P < 0.05). Images are maximum projection of acquired z-stack. Time shown as minutes:seconds. Time 0 represents anaphase onset. Vertical scale bars, 1 μm. Horizontal scale bar, 1min.

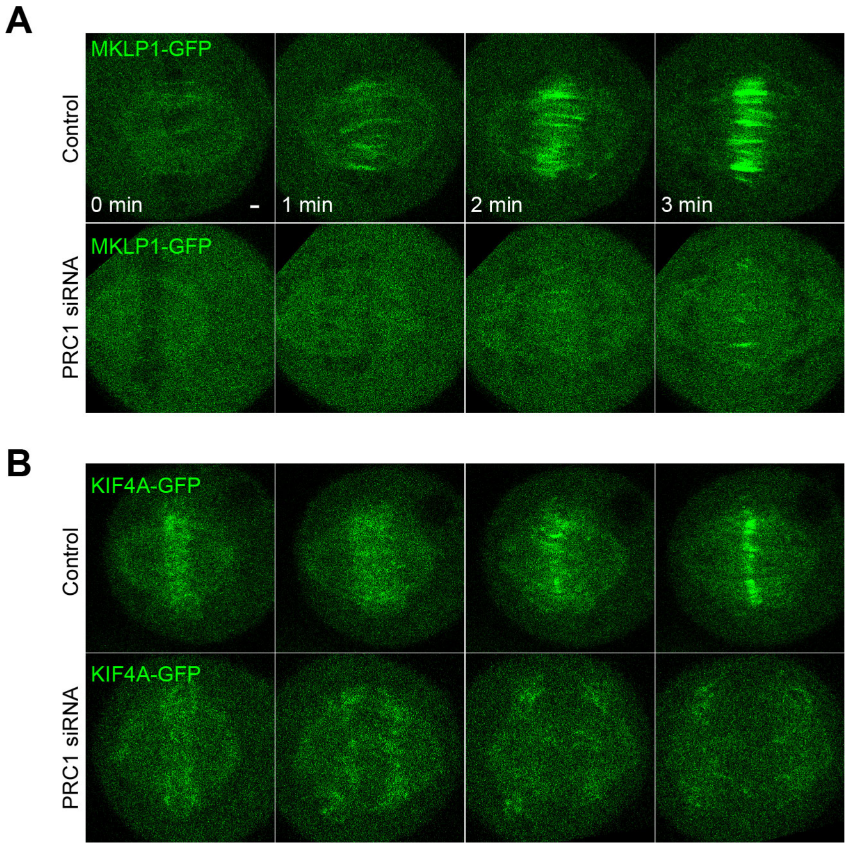


**Fig. S4.** Depletion of PRC1 removes its major binding partners, MKLP1 and KIF4A, from spindle midzone microtubules. (A) Live cell images of HeLA cell with stable expression of MKLP1-GFP from BAC in control and PRC1 siRNA-treated cell. (B) Live cell images of HeLA cell with stable expression of KIF4A-GFP from BAC in control and PRC1 siRNA-treated cell. Images are single z-planes. Time 0 represents anaphase onset. Scale bar, 1 μm.

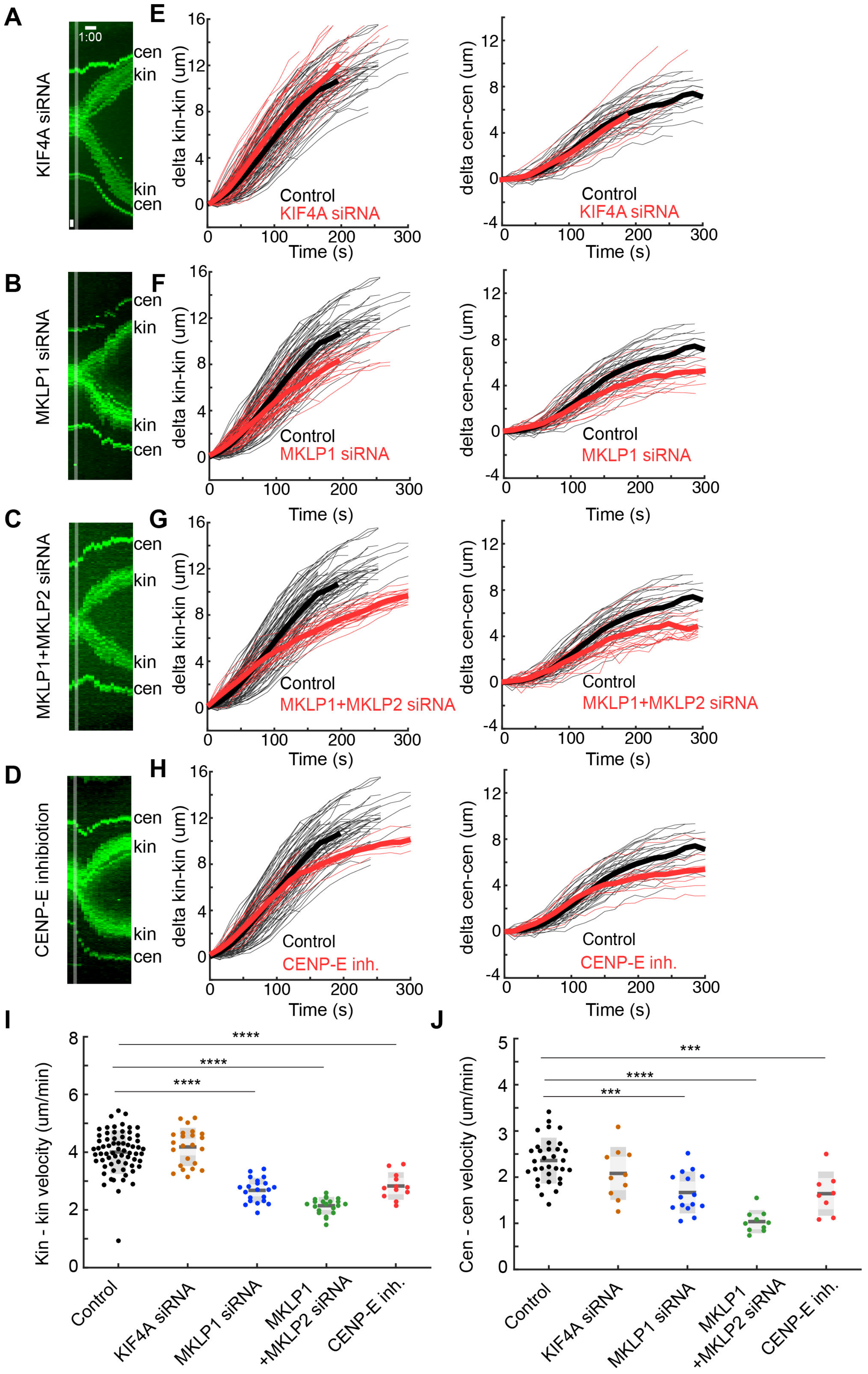
A
B
C
D
E
F
G
H
I
J
KIF4A siRNA
MKLP1 siRNA
MKLP1+MKLP2 siRNA
CENP-E inhibiotion
1:00
cen
kin
delta kin-kin (um)
delta cen-cen (um)
Time (s)
Control
KIF4A siRNA
MKLP1 siRNA
MKLP1+MKLP2 siRNA
CENP-E inh.
Kin - kin velocity (um/min)
Cen - cen velocity (um/min)
****
***
Control
KIF4A siRNA
MKLP1 siRNA
MKLP1
+MKLP2 siRNA
CENP-E inh.

**Fig. S5.** Depletion of PRC1 interacting partners has a minor effect anaphase spindle elongation. (A-D) Kymographs (consecutive maximal intensity projections onto the x-axis) of the live cell images of RPE-1 cells stably expressing CENP-A-GFP and centrin1-GFP in the siRNA depletions of KIF4A, MKLP1, MKLP1 + MKLP2 and CENP-E inhibition. (E-H) Plots of relative kinetochore and centrosome separation distance (delta) defined as the kinetochore-to-kinetochore (kin-kin) and centrosome-to-centrosome (cen-cen) distance at time t minus the kin-kin and cen-cen distance at t = 0, over time. Individual kinetochore and centrosome pairs (thin lines), mean (thick lines) for each condition from (A). Time 0 represents anaphase onset. (I, J) Quantification (univariate scatter plot) of velocity of separation of sister kinetochores (kin-kin) and spindle elongation (cen-cen) velocity. Boxes represent standard deviation (dark grey), 95% standard error of the mean (light grey) and mean value (black) for each condition from (A). Statistics: t test (***$P < 0.001$; ****$P < 0.0001$). Time shown as minutes:seconds. Vertical scale bars, 1 μm. Horizontal scale bar, 1min.

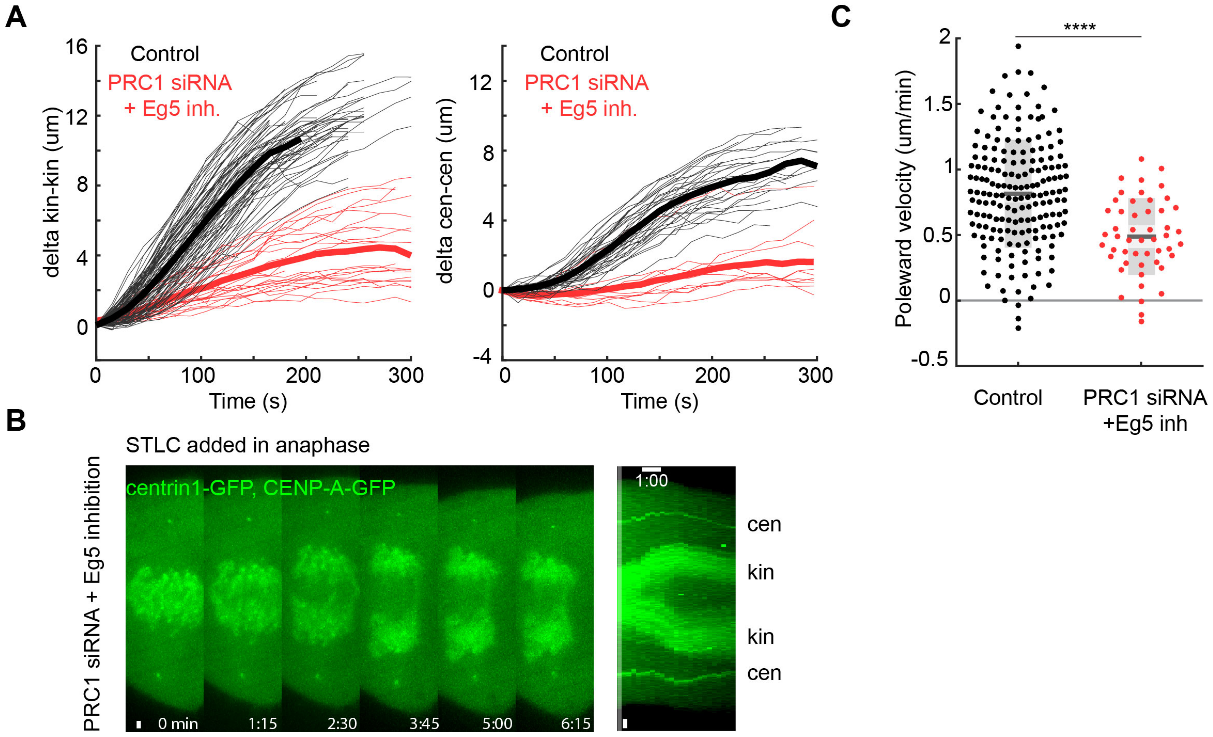


**Fig. S6.** Depletion of PRC1 combined with EG5 inhibition blocks spindle elongation in anaphase. (A) Plots of relative kinetochore and centrosome separation distance (delta) defined as the kinetochore-to-kinetochore (kin-kin) and centrosome-to-centrosome (cen-cen) distance at time t minus the kin-kin and cen-cen distance at t = 0, over time in RPE-1 cells stably expressing CENP-A-GFP and centrin1-GFP. Individual kinetochore and centrosome pairs (thin lines), mean (thick lines). Time 0 represents anaphase onset. (B) Live cell images of PRC1 siRNA depleted and S-trityl-L-cysteine (STLC)-treated RPE-1 cell show unperturbed spindle elongation. kin-kinetochore and cen-centrosome. Time shown as minutes:seconds. Time 0 represents STLC addition. (C) Quantification (univariate scatter plot) of kinetochore to pole velocities (poleward velocity). Boxes represent standard deviation (dark grey), 95% standard error of the mean (light grey) and mean value (black). Statistics: t test (****P < 0.0001). Images are maximum projection of acquired z-stack. Vertical scale bars, 1 µm. Horizontal scale bar, 1min.

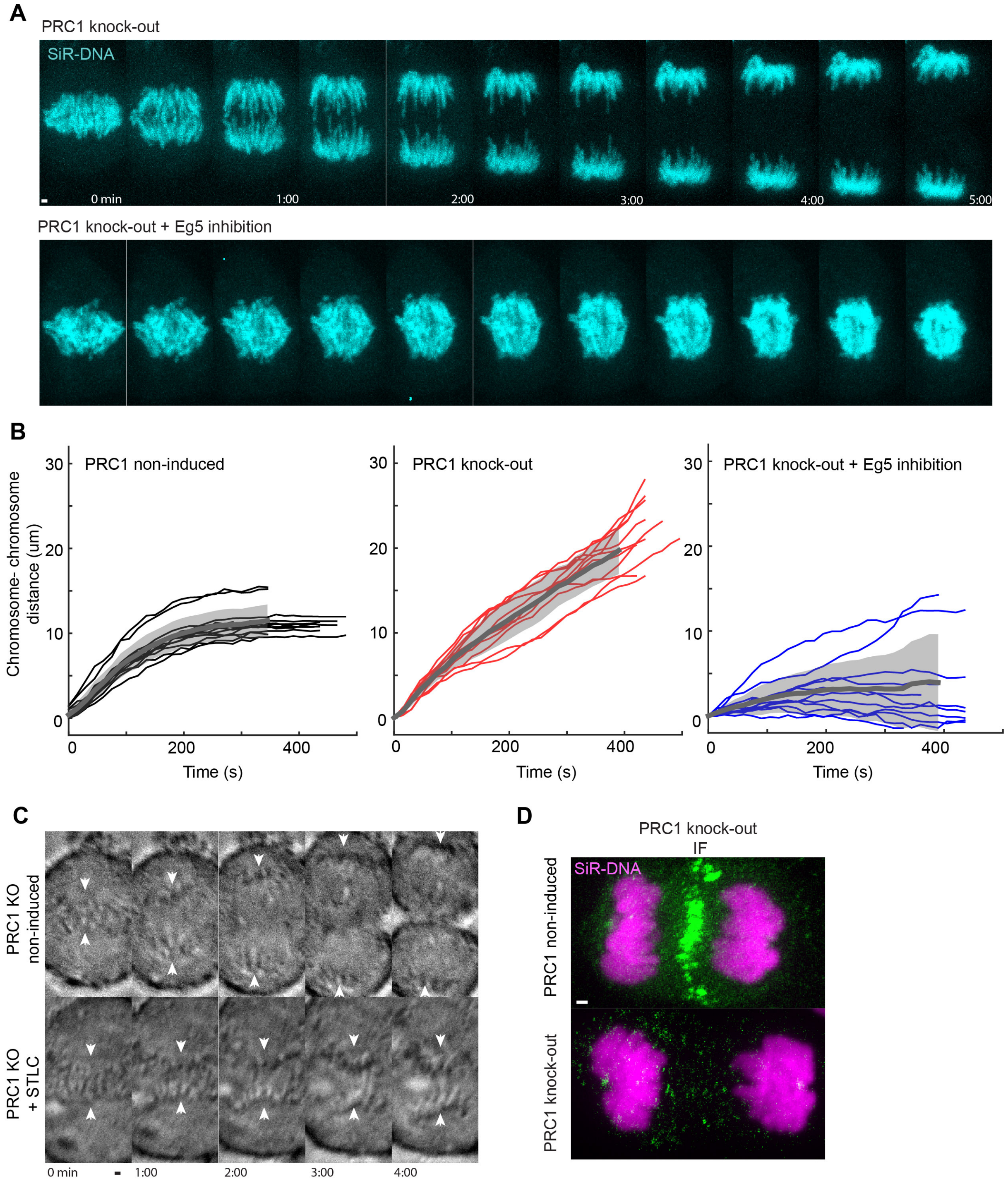


**Fig. S7.** PRC1 CRISPR knock-out (KO) combined with EG5 inhibition blocks spindle elongation in anaphase. (A) Live cell images of induced RPE-1 PRC1 CRISPR KO and PRC1 KO treated with STLC imaged 4 days after doxycycline induction. Silicon rhodamine (SiR)-DNA (cyan) was used for chromosome staining. (B) Plots of relative chromosome segregation distance (delta) defined as the chromosome-t-chromosome distance at time t minus the chromosome-chromosome distance at t = 0, over time. Individual chromosome pairs (thin lines), mean (thick grey lines) and SD (grey region) for indicated conditions. (C) Live cell brightfield images of induced RPE-1 PRC1 CRISPR KO and PRC1 CRISPR KO treated with STLC imaged 4 days after doxycycline induction. Note: chromosome segregation defects after STLC treatment are similar to ones observed in the same treatment but with chromosomes labelled with SiR-DNA. White arrowheads indicate position of chromosomes. (D) Immunofluorescence (IF) images of fixed non-induced PRC1 CRISPR KO and PRC1 CRISPR induced KO RPE-1 cells stained with AlexaFluor594 conjugated with PRC1 antibody (green) and silicon rhodamine (SiR)-DNA (magenta). Images are maximum projection of acquired z-stack. Time shown as minutes:seconds. Time 0 represents anaphase onset. Scale bars, 1 μm.

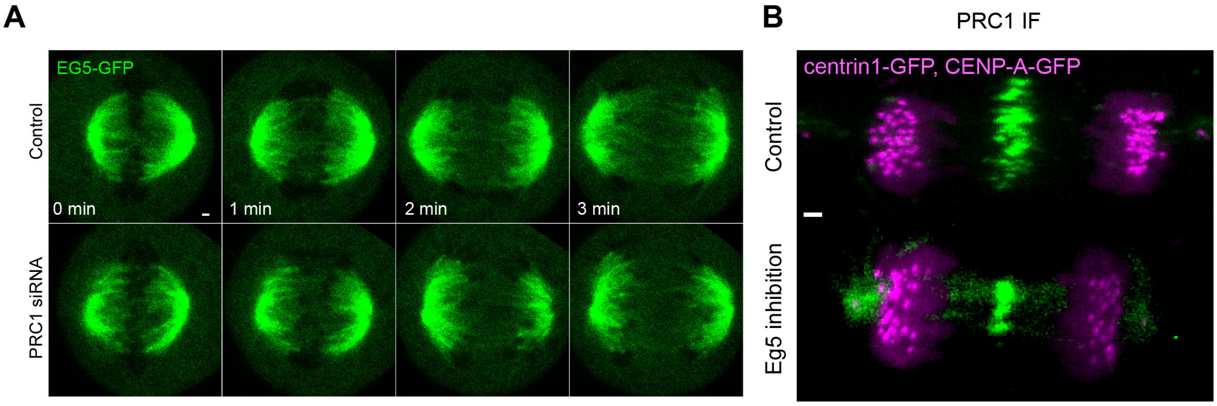


**Fig. S8.** Depletion of PRC1 does not affect EG5 localization on the spindle and *vice versa*. (A) Live cell images of control and PRC1 siRNA depleted HeLa cells stably expressing EG5-GFP from a BAC. Images are single z-planes. (B) Imunofluorescence (IF) images of fixed control and 10 min STLC-treated RPE-1 cells stably expressing CENP-A-GFP and centrin1-GFP (magenta) stained with AlexaFluor594 conjugated with PRC1 antibody (green). Images are maximum projection of acquired z-stack. Time 0 represents anaphase onset. Scale bars, 1 μm.

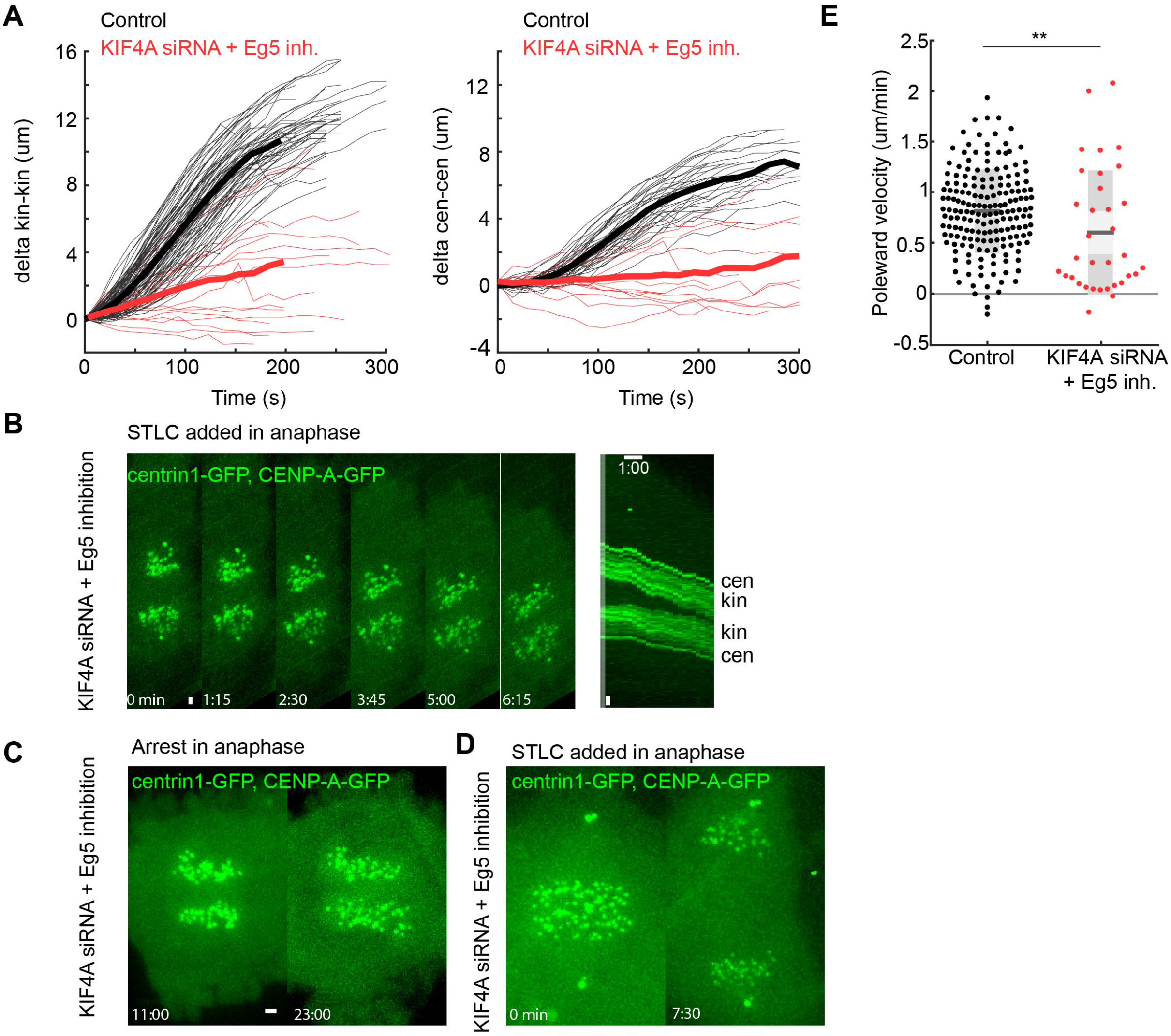


**Fig. S9.** Depletion of KIF4A combined with EG5 inhibition blocks spindle elongation in anaphase. (A) Plots of relative kinetochore and centrosome separation distance (delta) defined as the kinetochore-to-kinetochore (kin-kin) and centrosome-to-centrosome (cen-cen) distance at time t minus the kin-kin and cen-cen distance at t = 0, over time in KIF4A siRNA depleted and S-trityl-L-cysteine (STLC)-treated RPE-1 cells stably expressing CENP-A-GFP and centrin1-GFP. Individual kinetochore and centrosome pairs (thin lines), mean (thick lines). (B) Live cell images of a cell from (A) show perturbed spindle elongation when STLC is added in anaphase. kin-kinetochore and cen-centrosome. (C) Live cell images of KIF4A siRNA depleted and STLC-treated RPE-1 cell show long-term anaphase arrest phenotype when STLC is added in anaphase. (D) Live cell images of KIF4A siRNA depleted and STLC-treated RPE-1 cell, with addition of STLC in anaphase, show small rate of spindle elongation. Note: in few examples, if STLC was added in early anaphase, spindle elongation could be observed but to lesser degree when compared to controls. Time 0 represents start of STLC treatment. (E) Quantification (univariate scatter plot) of kinetochore to pole velocities (poleward velocity). Boxes represent standard deviation (dark grey), 95% standard error of the mean (light grey) and mean value (black). Statistics: t test (** $P < 0.01$). Images are maximum projection of acquired z-stack. Time shown as minutes:seconds. Vertical scale bars, 1 μm. Horizontal scale bar, 1min.

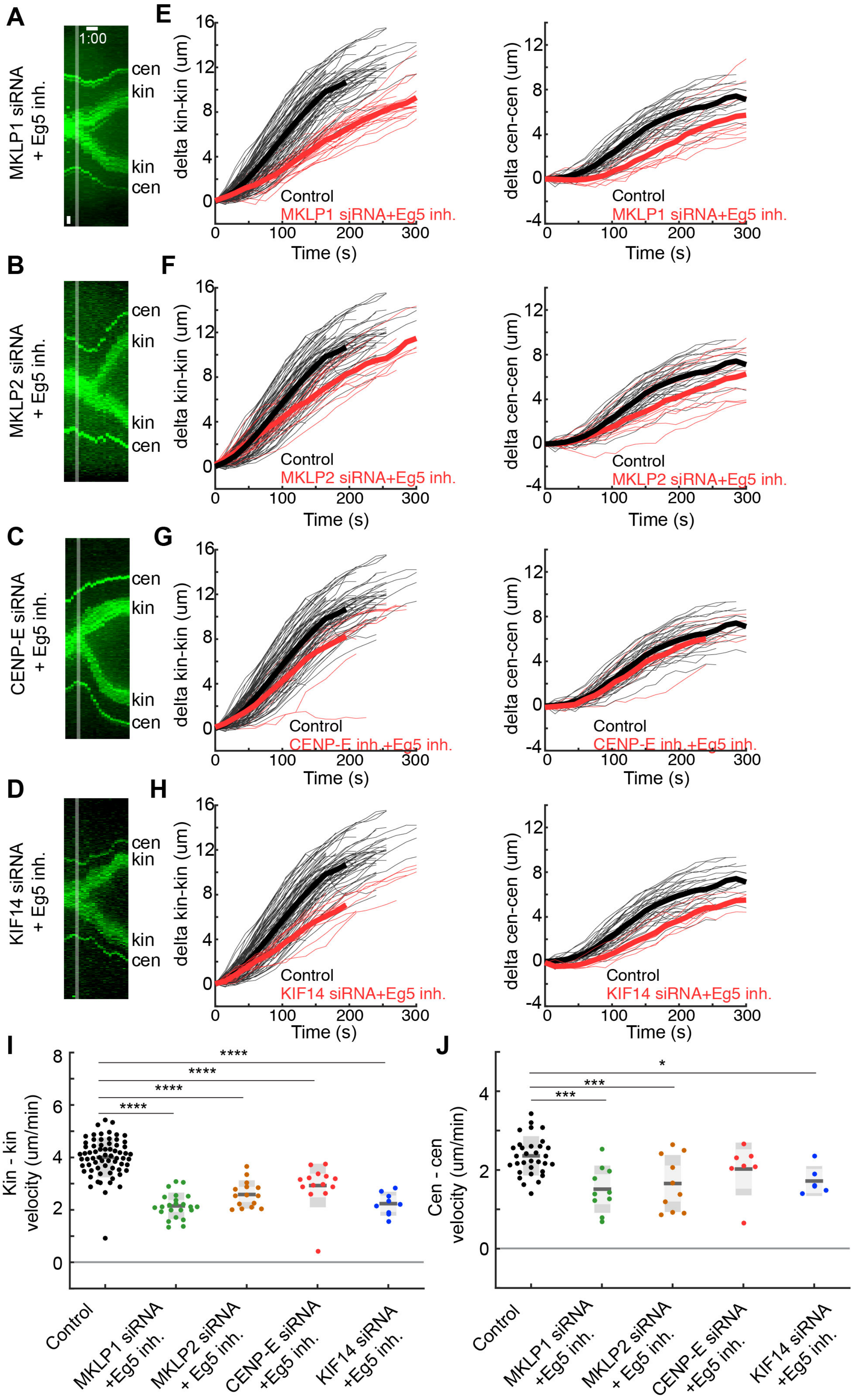
A
B
C
D
E
F
G
H
I
J
MKLP1 siRNA + Eg5 inh.
MKLP2 siRNA + Eg5 inh.
CENP-E siRNA + Eg5 inh.
KIF14 siRNA + Eg5 inh.
1:00
cen
kin
kin
cen
delta kin-kin (um)
delta cen-cen (um)
Time (s)
Control
MKLP1 siRNA+Eg5 inh.
MKLP2 siRNA+Eg5 inh.
CENP-E inh.+Eg5 inh.
KIF14 siRNA+Eg5 inh.
Kin - kin velocity (um/min)
Cen - cen velocity (um/min)
****
***
*
Control
MKLP1 siRNA +Eg5 inh.
MKLP2 siRNA + Eg5 inh.
CENP-E siRNA +Eg5 inh.
KIF14 siRNA +Eg5 inh.

**Fig. S10.** Depletion of PRC1 interacting partners with inactivation of EG5 has a minor effect on anaphase spindle elongation. (A-D) Kymographs (consecutive maximal intensity projections onto the x-axis) of the live cell images of RPE-1 cells stably expressing CENP-A-GFP and centrin1-GFP in the siRNA depletions of KIF4A, MKLP1, MKLP1 + MKLP2 and CENP-E inhibition combined with S-trityl-L-cysteine (STLC)-treatment. (E-H) Plots of relative kinetochore and centrosome separation distance (delta) defined as the kinetochore-to-kinetochore (kin-kin) and centrosome-to-centrosome (cen-cen) distance at time t minus the kin-kin and cen-cen distance at t = 0, over time. Individual kinetochore and centrosome pairs (thin lines), mean (thick lines) for each condition from (A). Time 0 represents anaphase onset. (I, J) Quantification (univariate scatter plot) of velocity of separation of sister kinetochores (kin-kin) and spindle elongation (cen-cen) velocity. Boxes represent standard deviation (dark grey), 95% standard error of the mean (light grey) and mean value (black) for each condition from (A). Statistics: t test (***$P < 0.001$; ****$P < 0.0001$). Vertical scale bars, 1 μm. Horizontal scale bar, 1min.

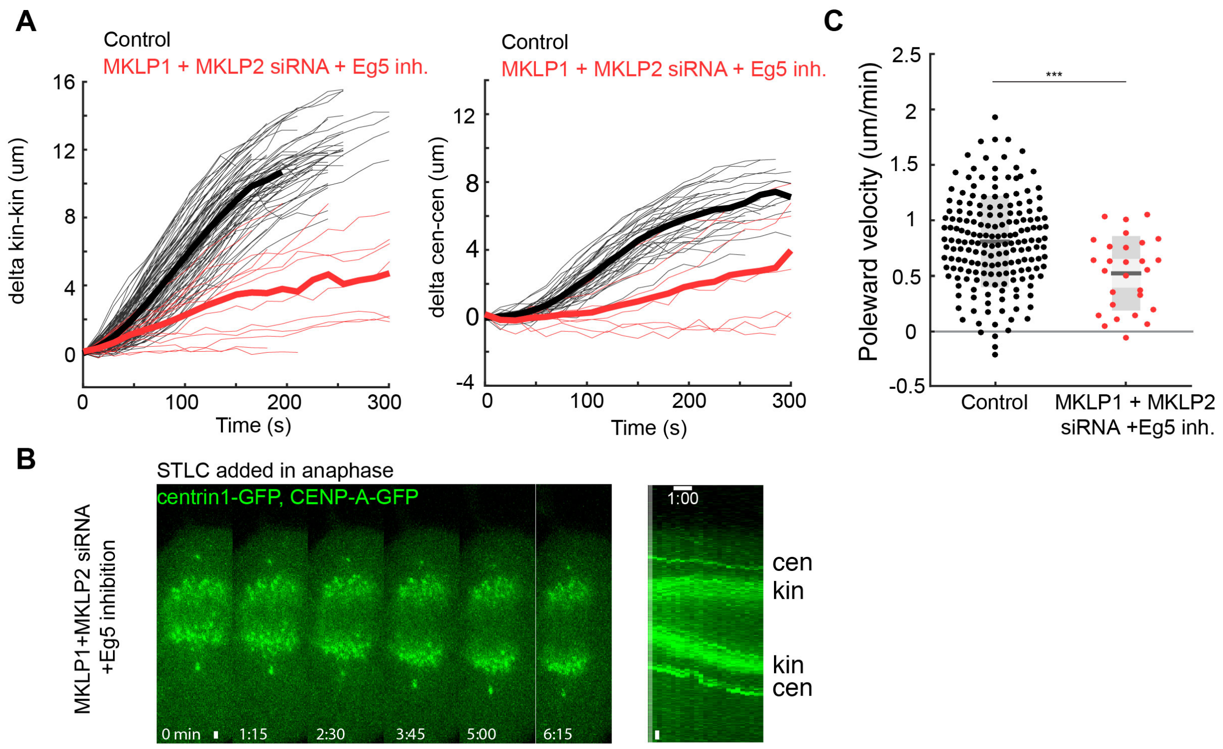


**Fig. S11.** Depletion of both kinesins-6 with inactivation of EG5 induces anaphase arrest. (A) Plots of relative kinetochore and centrosome separation distance (delta) defined as the kinetochore-to-kinetochore (kin-kin) and centrosome-to-centrosome (cen-cen) distance at time t minus the kin-kin and cen-cen distance at t = 0, over time in MKLP1 and MKLP2 siRNA depleted and S-trityl-L-cysteine (STLC)-treated RPE-1 cells stably expressing CENP-A-GFP and centrin1-GFP. Individual kinetochore and centrosome pairs (thin lines), mean (thick lines). (B) Live cell images of a cell from (A) show blocked spindle elongation. kin-kinetochore and cen-centrosome. Time 0 represents start of STLC treatment. Time 0 represents anaphase onset. (C) Quantification (univariate scatter plot) of kinetochore to pole velocities (poleward velocity). Boxes represent standard deviation (dark grey), 95% standard error of the mean (light grey) and mean value (black). Statistics: t test (***P < 0.001). Images are maximum projection of acquired z-stack. Time shown as minutes:seconds. Vertical scale bars, 1 μm. Horizontal scale bar, 1min.

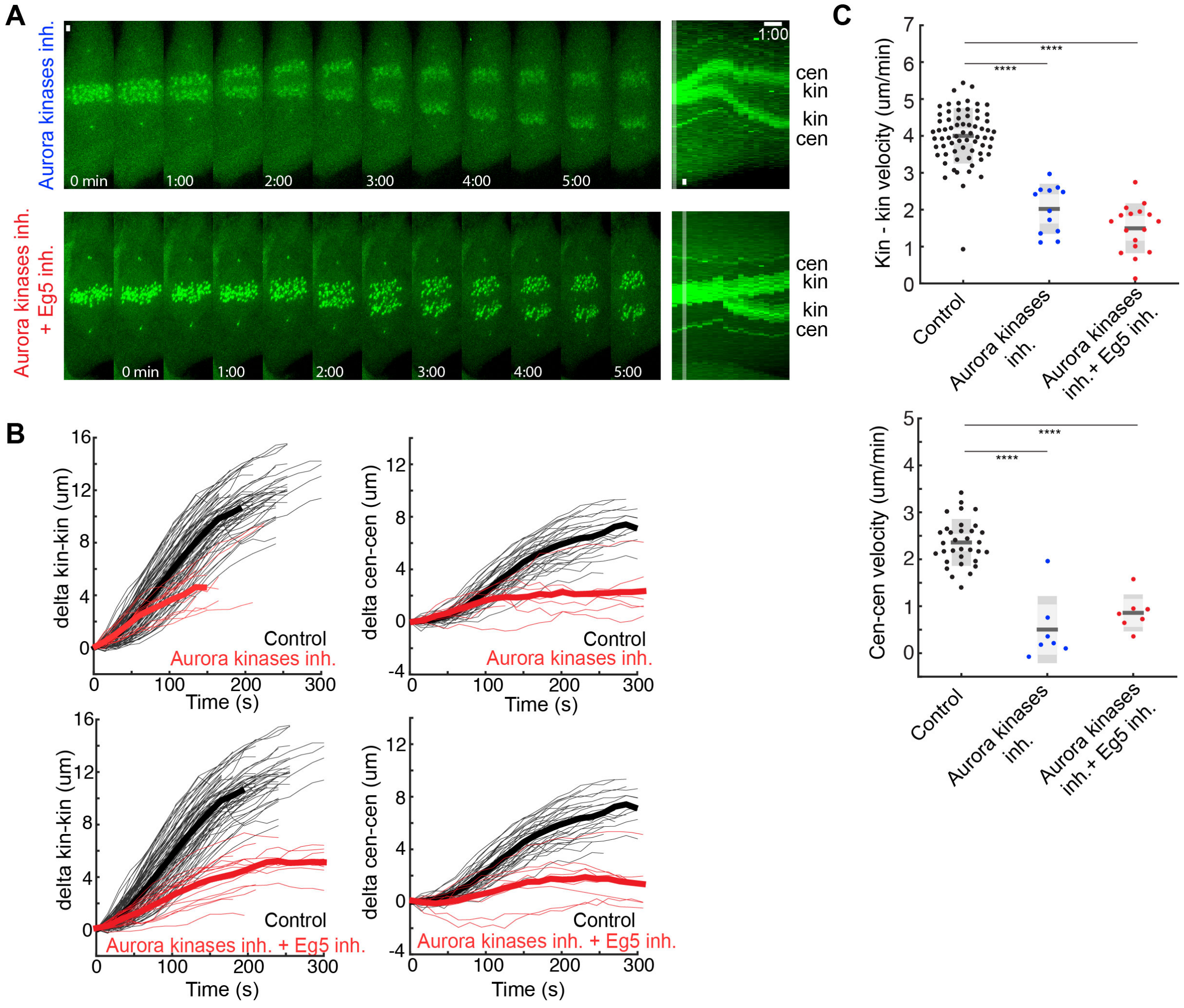


**Fig. S12.** Inhibition of Aurora kinases blocks spindle elongation. (A) Live cell images of RPE-1 cells stably expressing CENP-A-GFP and centrin1-GFP treated with ZM-447439 and co-treated with ZM-447439 and STLC that show blocked spindle elongation. kin-kinetochore and cen-centrosome. Time 0 represents anaphase onset. (B) Plots of relative kinetochore and centrosome separation distance (delta) defined as the kinetochore-to-kinetochore (kin-kin) and centrosome-to-centrosome (cen-cen) distance at time t minus the kin-kin and cen-cen distance at t = 0, over time in ZM-447439 treated and S-trityl-L-cysteine (STLC) co-treated RPE-1 cells. Individual kinetochore and centrosome pairs (thin lines), mean (thick lines). (C) Quantification (univariate scatter plot) of kinetochore to pole velocities (poleward velocity). Boxes represent standard deviation (dark grey), 95% standard error of the mean (light grey) and mean value (black). Statistics: t test (****P < 0.0001). Images are maximum projection of acquired z-stack. Time shown as minutes:seconds. Vertical scale bars, 1 μm. Horizontal scale bar, 1min.

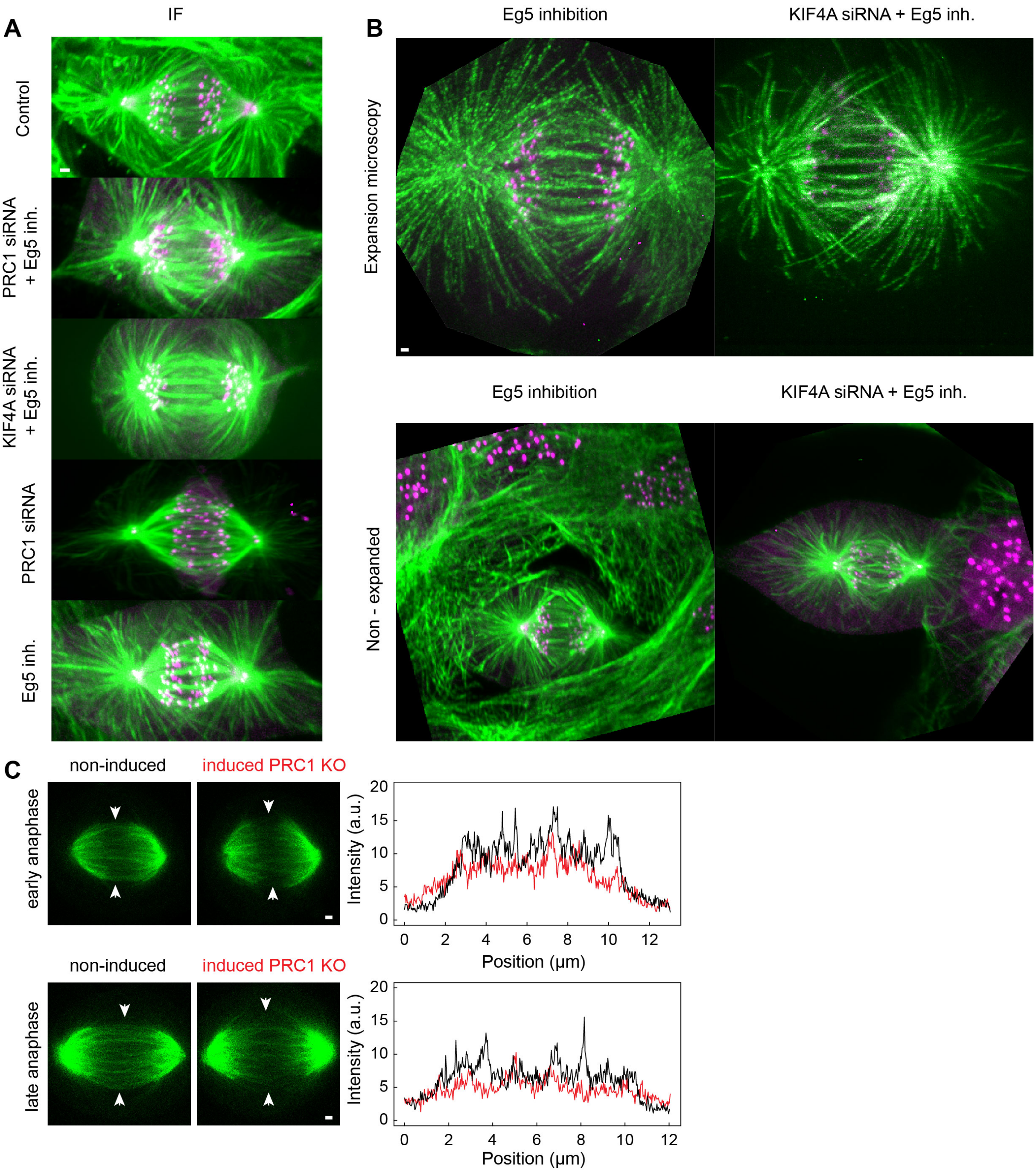


**Fig. S13.** Perturbation of spindle elongation is not due to the lack of microtubules in spindle midzone during anaphase. (A) Immunofluorescence (IF) images of fixed RPE-1 cells stably expressing CENP-A-GFP and centrin1-GFP (magenta) stained with AlexaFluor594 conjugated with α -tubulin antibody in indicated conditions. (B) Expanded and non-expanded images of fixed STLC-treated and KIF4A siRNA-depleted and STLC-treated RPE-1 cells stably expressing CENP-A-GFP and centrin1-GFP (magenta) stained with AlexaFluor594 conjugated with α-tubulin antibody (green). Expansion factor is estimated from measured spindle length to be 2.3x. Images are maximum projection of acquired z-stack. (C) STED images (single z-plane) of anaphase spindles in a live non-induced control CRISPR PRC1 knock-out (KO) and doxycycline induced PRC1 KO RPE-1 cells. Microtubules are labelled with SiR-tubulin. Arrowheads point to the spindle midzone region. Signal intensity profile of SiR-tubulin measured perpendicular to the spindle midzone. Note: signal intensities after

PRC1 KO are slightly decreased and bundles (signal intensity peaks) are less defined when compared to the non-induced control cells. Scale bars, 1 μm.

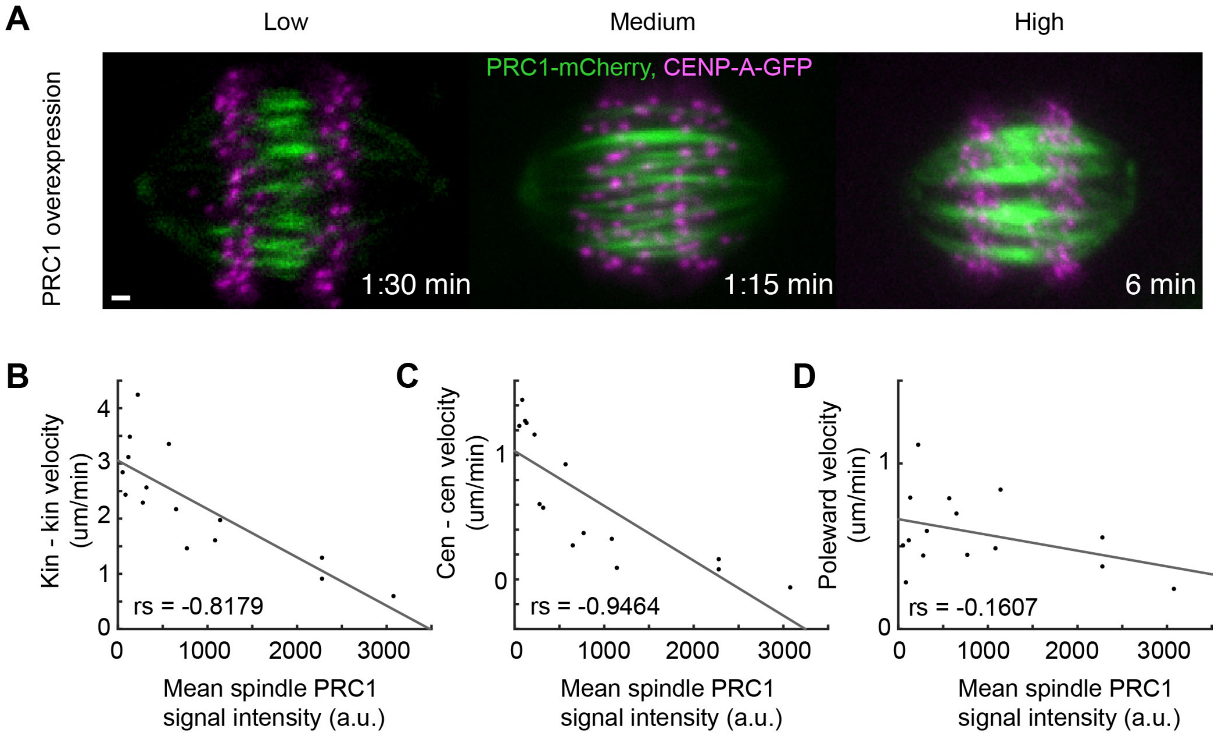


**Fig. S14.** PRC1 overexpression decreases spindle elongation velocities. (A) Live cell images of U2OS cells stably expressing CENP-A-GFP and transiently expressing various levels of PRC1-mCherry as indicated. (B-D) Linear regression and distribution of kinetochore separation (kin-kin), centrosome separation (cen-cen) and kinetochore-to-centrosome (poleward) velocity versus mean PRC1 signal intensity. rs, Spearman correlation coefficient, $P < 0.001$. Images are maximum projection of acquired z-stack. Time 0 represents anaphase onset. Time shown as minutes:seconds. Scale bar, 1 μm.

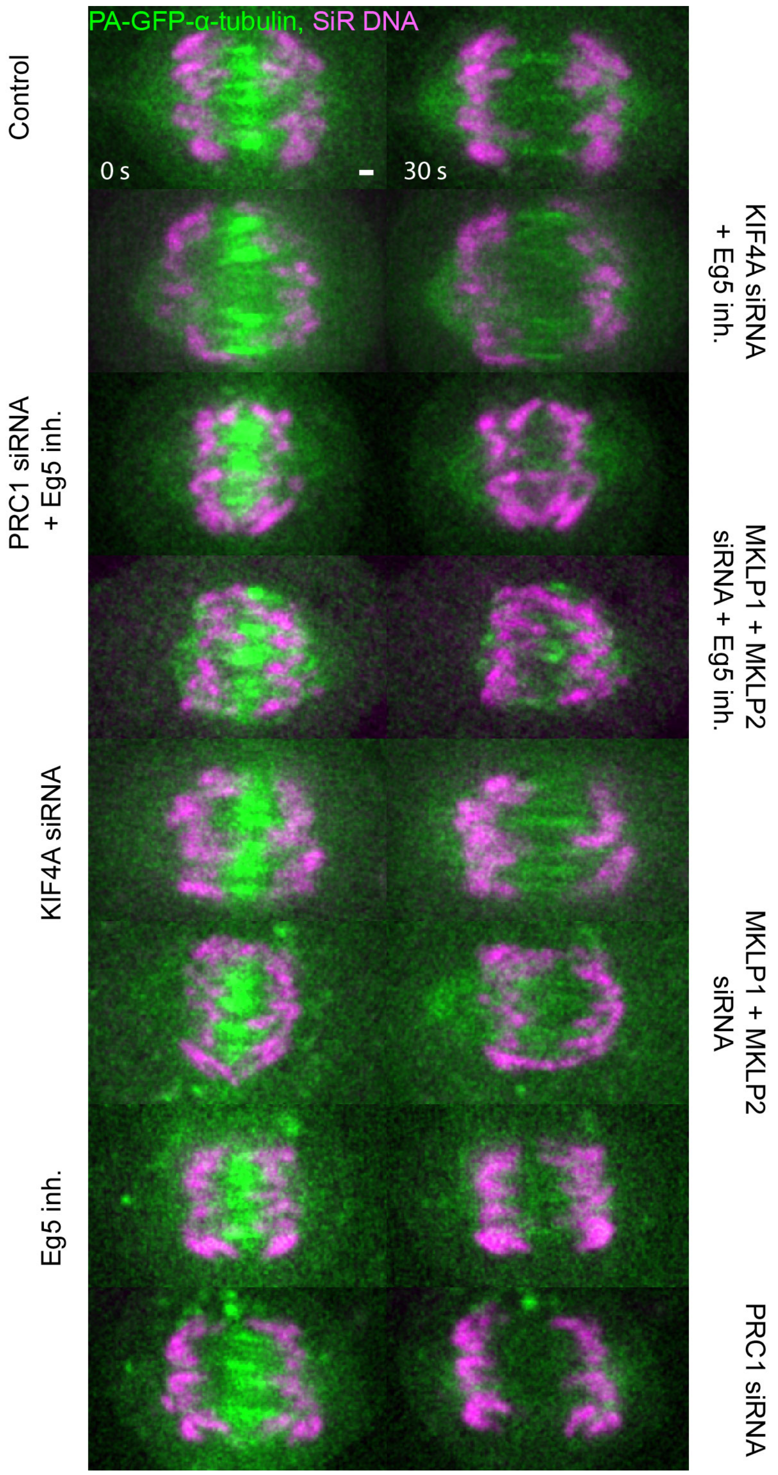


**Fig. S15.** Characterization of midzone stability in RPE-1 cells after different treatments using midzone photoactivation assay. Smoothed live cell images of RPE-1 cells after photoactivation of photoactivatable (PA)-GFP-α-tubulin in indicated conditions. Silicon rhodamine (SiR)-DNA (magenta) was used for chromosome staining. Time 0 represents photoactivation onset. Images are single z-planes. Time shown in seconds (s). Scale bars, 1 μm.

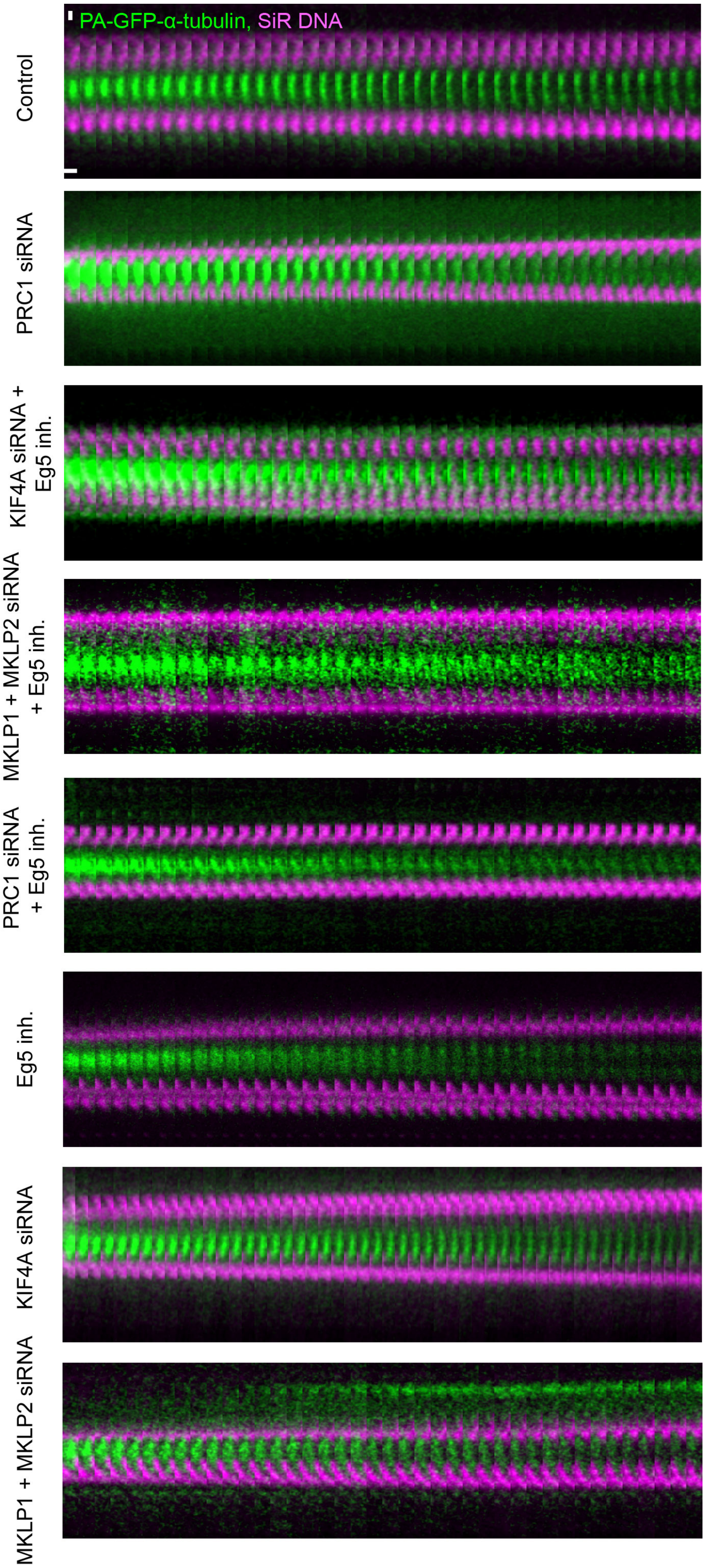
PA-GFP-α-tubulin, SiR DNA
Control
PRC1 siRNA
KIF4A siRNA + Eg5 inh.
MKLP1 + MKLP2 siRNA + Eg5 inh.
PRC1 siRNA + Eg5 inh.
Eg5 inh.
KIF4A siRNA
MKLP1 + MKLP2 siRNA

**Fig. S16.** Characterization of sliding in RPE-1 cells after different treatments using midzone photoactivation assay. Smoothed time-lapse live images of mitotic spindle midzone region in RPE-1 cells after photoactivation of photoactivatable (PA)-GFP-α-tubulin in indicated treatments. Silicon rhodamine (SiR)-DNA (magenta) was used for chromosome staining. Horizontal scale bar, 0.8s. Vertical scale bar, 1 μm.

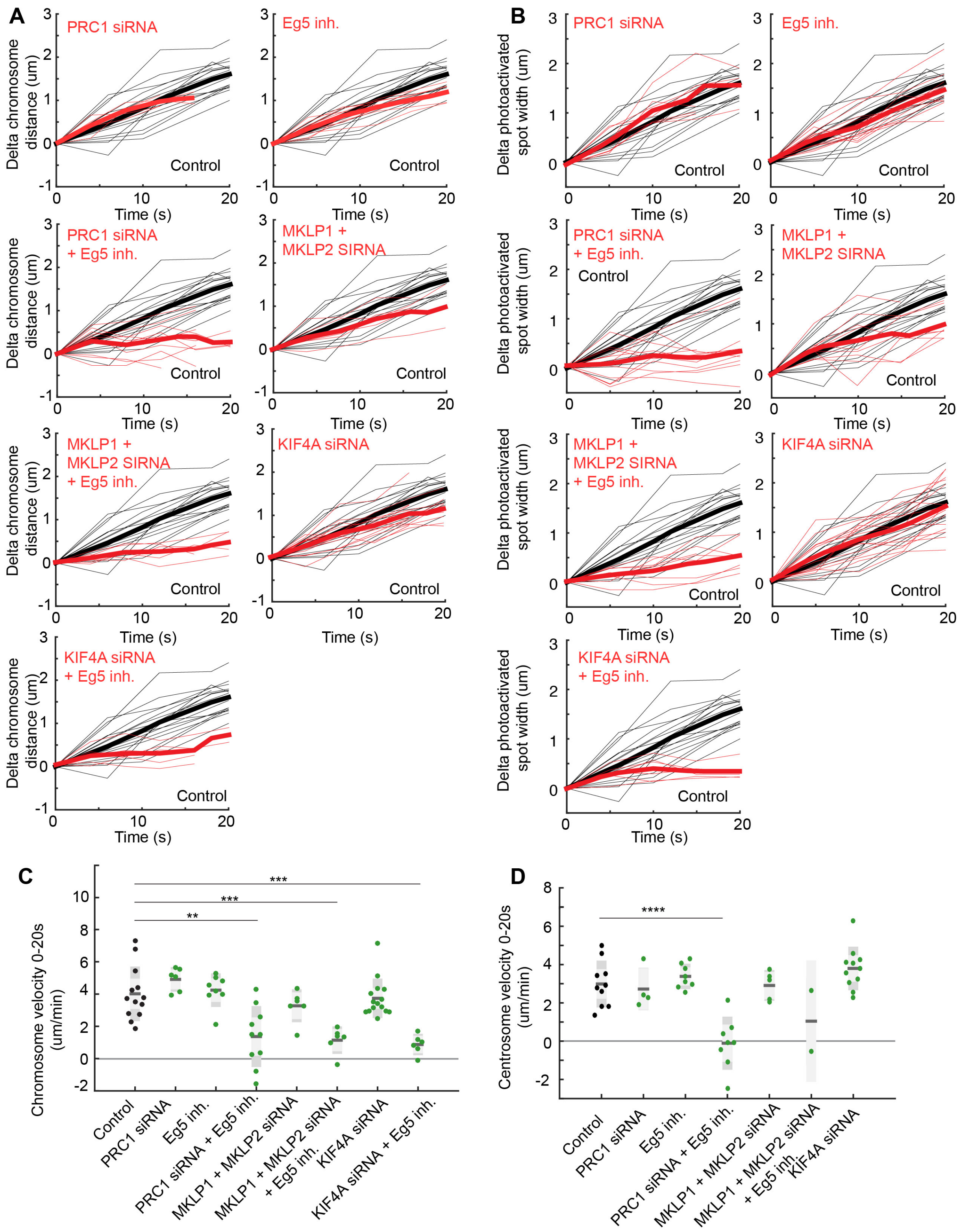


**Fig. S17.** Sliding of midzone microtubules and anaphase velocities are impaired after EG5 inhibition when combined with conditions perturbing KIF4A motor protein. (A) Plots of relative chromosome segregation distance (delta) defined as the chromosome-to-chromosome distance at time t minus the chromosome-to-chromosome distance at t = 0, over time in RPE-1 cells stably expressing CENP-A-GFP and centrin1-GFP in indicated conditions. Individual kinetochore and centrosome pairs (thin lines), mean (thick lines). (B) Plots of relative photoactivated spot length (delta) defined as the photoactivated spot length at time t minus the photoactivated spot length at t = 0, over time in RPE-1 cells in indicated conditions. Time 0 represents photoactivation onset. (C, D) Quantification (univariate scatter plot) of velocity of chromosome segregation and centrosome separation. Boxes represent standard deviation (dark grey), 95% standard error of the mean (light grey) and mean value (black) for indicated conditions. Statistics: t test (** $P < 0.01$; ***$P < 0.001$; ****$P < 0.0001$).
